# Co-expression patterns define epigenetic regulators associated with neurological dysfunction

**DOI:** 10.1101/219097

**Authors:** Leandros Boukas, James M. Havrilla, Aaron R. Quinlan, Hans T. Bjornsson, Kasper D. Hansen

## Abstract

Coding variants in genes encoding for epigenetic regulators are an emerging cause of neurological dysfunction and cancer. However, a systematic effort to identify disease candidates within the human epigenetic machinery (EM) has not been performed, and it is unclear whether features exist that distinguish between variation-intolerant and variation-tolerant EM genes, and between EM genes associated with neurological dysfunction versus cancer. Here, we rigorously define a set of 295 human genes with a direct role in epigenetic regulation (writers, erasers, remodelers, readers). Systematic exploration of these genes reveals that while individual enzymatic functions are always mutually exclusive, readers often also exhibit enzymatic activity as well (dual function EM genes). We find that the majority of EM genes are very intolerant to loss-of-function variation, even when compared to the dosage sensitive group of transcription factors. Using this strategy, we identify 103 novel EM disease candidates. We show that the intolerance to loss-of-function variation is driven by the protein domains encoding the epigenetic function, strongly suggesting that disease is caused by a perturbed chromatin state. Unexpectedly, we also describe a large subset of EM genes that are co-expressed within multiple tissues. This subset is almost exclusively populated by extremely variation-intolerant EM genes, and shows enrichment for dual function EM genes. It is also highly enriched for genes associated with neurological dysfunction, even when accounting for dosage sensitivity, but not for cancer-associated EM genes. These findings prioritize novel disease candidate EM genes, and suggest that the co-expression itself may play a functional role in normal neurological homeostasis.

## Introduction

The chromatin landscape of any cell is shaped and maintained by the epigenetic machinery (EM), hereafter defined as the group of proteins that can catalyze the addition or removal of epigenetic marks (writer or erasers, respectively), bind to preexisting marks (readers), or use the energy of ATP hydrolysis to alter the local chromatin environment via mechanisms such as nucleosome sliding (remodelers) (Fahrner and Bjornsson, 2014; Allis and Jenuwein, 2016). Recently, some EM genes have been associated with human diseases, with the most prevalent disease phenotypes falling broadly under the categories of neurological dysfunction (De Rubeis et al., 2014; McCarthy et al., 2014; Bjornsson, 2015; Singh et al., 2016; Deciphering Developmental Disorders Study, 2017), and cancer (Vogelstein et al., 2013; Garraway and Lander, 2013; Feinberg, Koldobskiy, and Göndör, 2016); those associations have indicated that the vast majority of known disease causing EM genes are haploinsufficient (Garraway and Lander, 2013; Bjornsson, 2015).

This paper addresses three main questions. First, how many additional disease candidate EM genes are there? Existing estimates (Khare et al., 2012; Medvedeva et al., 2015) suggest that EM genes with ascribed roles in disease only form a minority of the whole group. Thus, the number of additional disease candidates that a comprehensive EM gene list will harbor is unclear. It is also unknown whether disease genes tend to be evenly distributed among classes (e.g. erasers vs remodelers) and subclasses (e.g. histone methyltransferases vs histone acetyltransferases) of the machinery; such patterns could reflect the relative contribution of those categories to normal cellular function. Second, is the lost epigenetic function of these genes the most likely cause of disease? Studies in model systems have indicated that the domains mediating the epigenetic function can be dispensable (Dorighi et al., 2017; Rickels et al., 2017). This raises the possibility that, even among known EM disease genes, the phenotype might have some alternative mechanistic basis. Third, are there expression signatures characteristic of disease candidates? In other words, are the expression patterns of EM genes that are intolerant to variation different from those of variation-tolerant EM genes? Related to this question, it would be of particular interest if there also exist expression signatures that distinguish between EM genes associated with neurological dysfunction versus those associated with cancer. Such signatures could not only prioritize candidate genes for specific phenotypes, but also provide insights into novel disease mechanisms.

To answer those questions, we perform a systematic investigation of the human epigenetic machinery with respect to its composition, its tolerance to variation, and its expression in a diverse set of tissues. We first define a set of 295 human EM genes on the basis of protein domain annotations (Hunter et al., 2009); this strategy has been very helpful in elucidating the general characteristics of Transcription Factors (TFs) (Vaquerizas et al., 2009). We then leverage large scale data on human genetic variation (Lek et al., 2016) to demonstrate that the most EM genes are highly intolerant to the loss of a single allele. We show that this intolerance is even more pronounced than for TF genes, and is predominantly attributable to the domains that mediate the epigenetic function. In total, we identify 103 dosage sensitive EM genes which currently have no existing disease association. Unexpectedly, we discover that more than a third of EM genes are co-expressed throughout multiple tissues. This co-expressed subset almost exclusively contains EM genes with severe dosage sensitivity, and is enriched for a unique class of EM genes that have both reading and enzymatic functions, nearly all of which are variation-intolerant. The co-expressed ensemble also shows enrichment for EM genes associated with neurological dysfunction (independently of dosage sensitivity), but not for those linked to cancer. These results strongly suggest that this phenomenon has functional relevance, particularly in the brain. Furthermore, they prioritize disease candidates, and distinguish those likely to lead to neurological dysfunction.

## Results

### The modular composition of the epigenetic machinery

We defined EM genes as genes whose protein products contain domains classifying them as chromatin remodelers, or as writers/erasers/readers of DNA or histone methylation, or histone acetylation. Then, we utilized the UniProt database (UniProt Consortium, 2015), combined with InterPro domain annotations (Hunter et al., 2009), to systematically compile a list of all such human genes (Methods; a full list of the domains used for classification is provided in Supplementary Table1). This stringent, domain-based definition minimizes the risk of false positives. We found a total of 295 EM genes (Figure 1a,b and Supplementary Table 2), the vast majority of which belong to the histone machinery (Figure 1a), and only a small fraction are remodelers or components of the DNA methylation machinery (Figure 1a). The two latter categories overlap the histone machinery; most remodelers are also readers of either histone methylation or acetylation, whereas the overlap between the DNA methylation and histone components is multifaceted (Methods).

**Figure 1.**
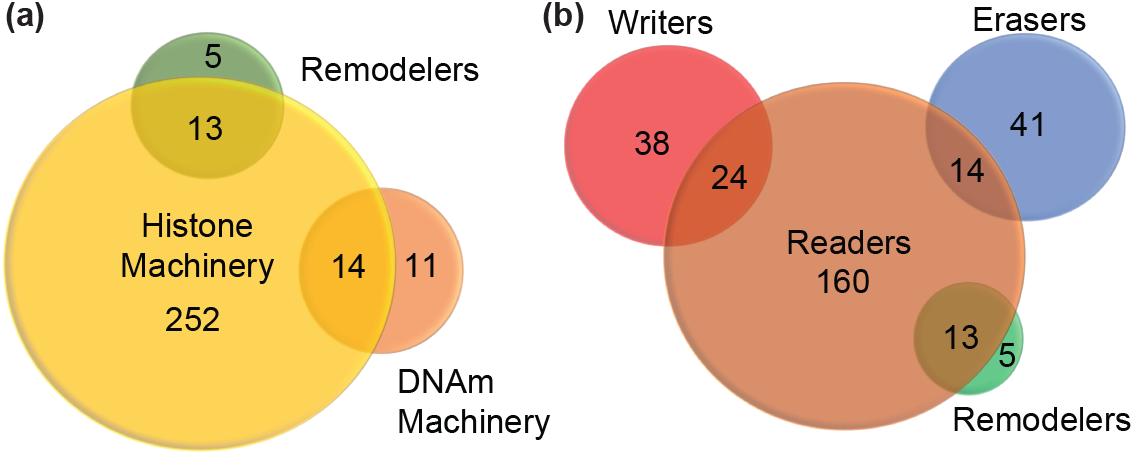
The modular composition of the epigenetic machinery. **(a)** Venn diagram illustrating the 3 broad categories of the epigenetic machinery (histone machinery, DNA methylation machinery, and remodelers), their relative sizes, and their mutual relationships. **(b)** Venn diagram illustrating the 4 broad “action” categories of the machinery (writers, erasers, remodelers, and readers), their relative sizes, and their mutual relationships. The modularity of this organization is evident, with some reader components exhibiting enzymatic functions and/or more than one reading functions. In contrast, the individual enzymatic component types are pairwise mutually exclusive.

Considering the categorization of EM genes into readers, writers, erasers and remodelers, we found that the readers comprise the biggest group (n = 211), and the remodelers the smallest (n = 18) (Figure 1b). The writer and eraser groups are comparable in size (n = 62, n = 54, respectively) (Figure 1b). We observed that the three enzymatic categories (writers, erasers, and remodelers) are pairwise mutually exclusive (Figure 1b). In contrast, we saw a subgroup of 52 genes encoding proteins which harbor both an enzymatic and a reader domain (Figure 1b), suggesting that these factors have dual epigenetic function; we will refer to these genes as dual function EM genes. These are not the only EM genes with a possible dual function: within the reader category there are 32 genes capable of recognizing more than one type of mark; we termed those dual readers, and find that some of them (n = 7) also have enzymatic activity. Furthermore, we observed that the same reading function can be mediated by different domains within a single protein; among the 178 readers of histone methylation we found 23 proteins which contain 2 distinct reading domains. We also observe that 9 of the domains defining EM genes can be present in multiple copies within the same gene, with the exact multiplicity ranging from 2 to 8 (calculated for the EM genes in Supplementary Table 4). Moreover, using a previously generated, high confidence list of 1,254 human transcription factors (Vaquerizas et al., 2009; Barrera et al., 2016), we found that 20 EM genes (12 of which are members of the PRDM family of histone methyltransferases), have a DNA binding domain found in transcription factors, suggesting their involvement in more than one aspect of transcriptional regulation.

### The human epigenetic machinery is highly intolerant to variation and contains many additional disease candidates

To identify novel EM disease candidates, we systematically investigated the tolerance of the entire EM group to loss-of-function variation. To achieve this, we used the ExAC database coupled with the pLI score (Lek et al., 2016), a metric which ranges between 0 and 1 and measures the extent to which a given gene tolerates heterozygous loss-of-function variants. In particular, genes with a pLI of more than 0.9 have been described as highly dosage sensitive (Lek et al., 2016), with virtually all known haploinsufficient human genes belonging to this category (Lek et al., 2016). A similar approach, focused only on the histone methylation machinery, was very recently used to derive candidate genes for developmental disorders (Faundes et al., 2018). In total, ExAC provides a pLI for 18,225 human genes, of which 281 are EM genes. First, we observed that EM genes have significantly higher pLI scores compared to all other genes (Wilcoxon rank sum test, *p* < 2.2 · 10^−16^, Figure 2a) and show substantial enrichment in the highly intolerant category (Fisher’s test, *p* < 2.2 · 10^−16^, odds ratio = 7.7). Given that pLI is a measure of haploinsufficiency, genes encoded on the X and Y chromosomes were not considered in this comparison (see Supplementary Methods for more details on EM genes encoded on the sex chromosomes).

**Figure 2.**
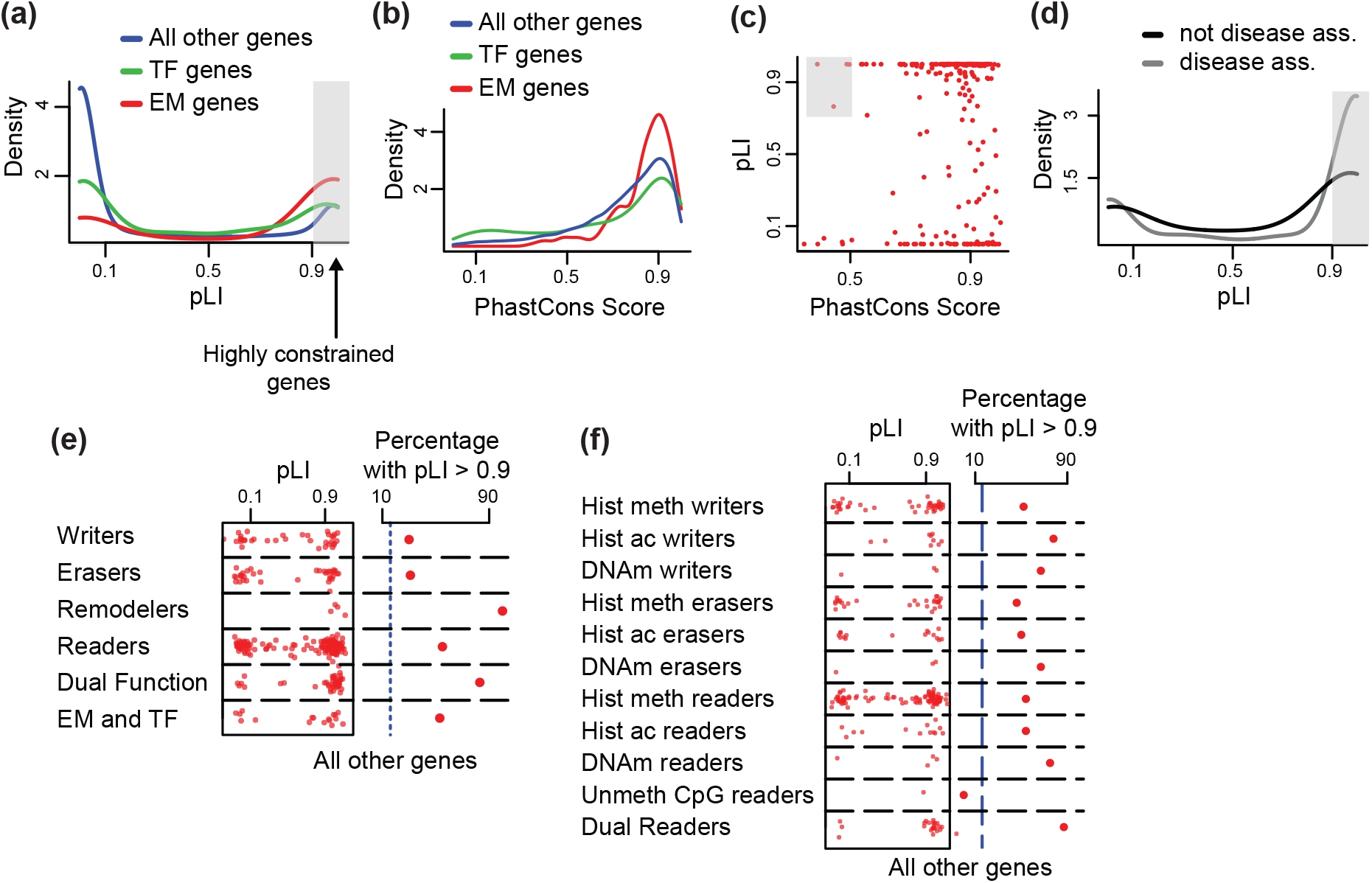
A large subset of components of the epigenetic machinery are very intolerant to variation. **(a)** The pLI distribution of EM genes (red curve) indicates their high enrichment in the pLI > 0.9 category (gray shaded area), which far exceeds that of TF genes (green curve), and that of all other genes (blue curve). **(b)** The PhastCons score distribution of EM genes (red curve) shows that they also exhibit higher across-species conservation than other genes (blue curve), and TF genes (green curve). **(c)** EM genes with high pLI also have a high PhastCons score. Four genes have PhastCons < 0.5 and high pLI (shaded area). **(d)** The pLI distribution of disease associated EM genes vs. non-disease associated EM genes. **(e)** Individual component classes display different levels of intolerance, with remodelers and dual function EM genes being the most intolerant. Each category exhibits higher dosage sensitivity than other human genes (blue dashed line). **(f)** Further dividing the parts of the machinery into subclasses provides a more detailed view of the mutational constraint of individual EM gene classes.

We next compared EM to TF genes; this is a natural comparison, since EM proteins are usually recruited to target sites by TFs (Lappalainen and Greally, 2017), and TF genes have been shown by previous analyses to be mostly haploinsufficient (Jimenez-Sanchez, Childs, and Valle, 2001; Seidman and Seidman, 2002). Using the 1,155 TF genes in ExAC, we first showed that they have significantly higher pLI compared to other genes (Wilcoxon rank sum test, *p* < 2.2 · 10^−16^, Figure 2a), although they are less dosage sensitive than previously suggested (Jimenez-Sanchez, Childs, and Valle, 2001), illustrating the value of our comprehensive approach in yielding accurate estimates. Comparing TF to EM genes however, we observed that EM genes have higher pLIs (Wilcoxon rank sum test, *p* < 2.2 · 10^−16^, Figure 2a) and are more strongly enriched in the highly dosage sensitive category (Fisher’s test, *p* < 2.2 · 10^−16^, odds ratio = 4.4). As an approach complementary to studying variation within the human population, we also examined the across-species conservation of EM genes, and obtained similar results (Figure 2b,c).

After splitting EM genes into those with existing disease associations and those with no reported link to disease (the latter constituting approximately 70%; Methods), we discovered that in both the disease and the non-disease associated groups there exist many EM genes with elevated pLI scores, (Figure 2d), although the disease associated ones exhibit higher skewing. It is notable that EM genes which are only associated with cancer have high pLIs (median pLI = 0.98, percentage with pLI> 0.9 = 65%). There is no *a priori* reason to expect this for somatic cancer driver genes, as pLI scores were derived after only excluding individuals with severe pediatric disease (Lek et al., 2016). Overall, this result strongly suggests the existence of additional EM disease genes. Among 163 EM genes in ExAC with no reported link to disease, 79 have a pLI greater than 0.9. Additionally, there are 24 EM genes that have only been associated with cancer but whose pLI is greater than 0.9, suggesting that they also cause some other phenotype. In total, this leads to 103 novel EM disease candidates (Supplementary Table 3).

### Dual function EM genes and remodelers are the most variation-intolerant categories

We next explored the loss-of-function variation tolerance of the different types of machinery components. Chromatin remodelers are an extremely intolerant group, whereas both the writers and the erasers are equally distributed among the high and low pLI groups (Figure 2e). Collectively, the three enzymatic EM classes show surprisingly high mutational constraint (Fisher’s test, *p* = 9.82 · 10^−9^, odds ratio = 4.1 for enrichment of EM genes with enzymatic but not reading function in the pLI > 0.9 category). Readers are more skewed towards the high pLI category than writers and erasers (Figure 2e). Despite the differences between these single-function classes, dual function EM genes are extremely constrained, regardless of the specific enzymatic function; this underscores the importance of this unique category (Figure 2e). Although histone methyltransferases (HMTs) appear less constrained than histone acetyltransferases (HATs) (Figure 2f), it is worth pointing out that all but one of the HMTs also have a reader domain, and there is no difference in variation tolerance between dual-function HMTs and dual-function HATs (Wilcoxon rank sum test, *p* = 0.61). Two out of the three DNA methyltransferases (*DNMT1* and *DNMT3B*) are constrained (Figure 2f), as is the case for DNA demethylases (with *TET2* being the only tolerant member, Figure 2f), whereas histone demethylases and deacetylases are approximately evenly divided in the high and low pLI categories (Figure 2f). This analysis also highlighted dual readers as a very intolerant group (Figure 2f). Finally, we note that all genes encoding for *CxxC* domain proteins, which recognize unmethylated CpG dinucleotides (Lee, Voo, and Skalnik, 2001), show very high dosage sensitivity (median pLI = 0.97), with three of them being dual readers.

### The intolerance to variation is primarily driven by the domains mediating the epigenetic function

A recent study showed that *Drosophila* embryos with a catalytically inactive version of *trr* (a homolog of the mammalian histone methyltransferases *KMT2C* and *KMT2D*) develop normally, despite altered histone methylation patterns (Rickels et al., 2017). This example shows that in some cases the inactivation of an epigenetic domain (in this case, the SET domain) might not have severe, easily detectable consequences. Our previous analysis is unable to determine if the observed variation intolerance is driven by the presence of epigenetic, or other non-epigenetic domains. We therefore asked if the EM-specific domain(s) in an EM gene had a different local mutational constraint than other domains in the same gene. To answer this, we used the constrained coding region (CCR) model (Havrilla et al., 2017) to derive a constraint score at the domain level, which we call CCR local constraint. This score reflects how devoid the domain is of missense or loss of function mutations in the gnomAD database (Lek et al., 2016), compared to other similar regions. It ranges from 0 to 100, with values close to 100 indicating highly constrained domains (Methods). We were able to score 237 out of 295 EM genes.

Under the hypothesis that EM-specific domains are not contributing to the observed variation intolerance of EM genes, there should be no difference in CCR local constraint between EM-specific domains of high pLI and low pLI EM genes. In contrast, we found that EM-specific domains of high pLI genes (greater than 0.9 pLI) have higher CCR local constraint than those of low pLi genes (less than 0.1 pLI) (Figure 3a). To explore this in more detail, we compared the within-gene average CCR local constraint of EM-specific domains to that of non EM-specific domains in EM genes. We observed that EM-specific domains have higher CCR local constraint than non EM-specific domains, but, as expected, only for high pLI EM genes (paired t-test, *p* = 0.009, Figure 3b). In this analysis we included 88 out of 142 EM genes with high pLI, which had 1 or more non EM-specific domains. Excluded were 54 high pLI EM genes without a non EM-specific domain; for these genes the EM domain is the only domain which can drive pLI. While EM-specific domains in general have higher CCR local constraint, there are 32 high pLI genes where this is not true. These 32 genes include all members of the PRDM family with high pLI, in which the highest CCR local constraint is observed in their C2H2 Zinc Finger domain. Our approach is however conservative, as there are domains that do not contribute catalytic or reading activity (and are thus labeled as non EM-specific), but are nevertheless found only in EM genes (see Methods). Returning to our initial example, in *KMT2D* the highest constraint is seen in the post-SET domain, whose precise functional role is not well understood, but it is not encountered outside of EM genes.

**Figure 3.**
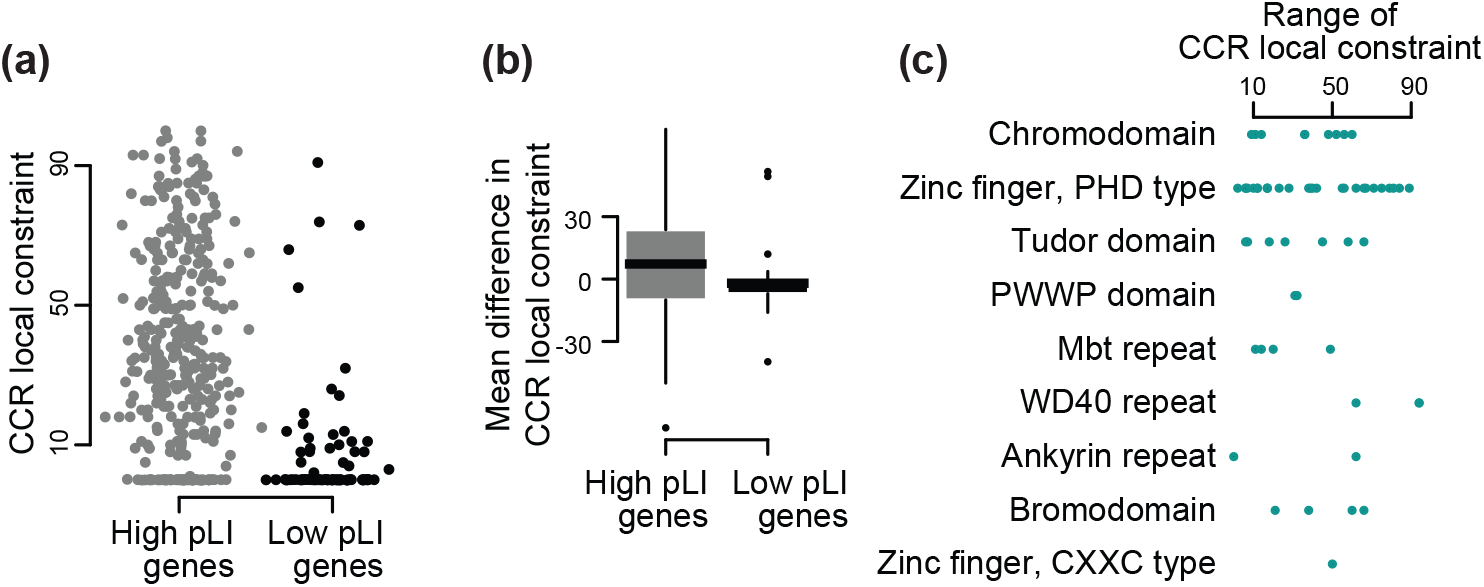
The protein domains known to mediate epigenetic function drive the observed constraint of EM genes. **(a)** The EM-specific protein domains of high pLI (> 0.9) EM genes are more highly constrained than those of low pLI (< 0.1) EM genes. **(b)** The distribution of within-gene differences in the average CCR local constraint of EM-specific domains minus the average CCR local constraint of non EM-specific domains shows that EM-specific domains are more intolerant to variation. As a control, for low pLI EM genes, no substantial difference in local constraint between EM-specific and non EM-specific domains is observed. **(c)** EM reader domains that appear in more than 1 copy within the same gene show within-gene variability in CCR local constraint (each point corresponds to the range of CCR local constraint scores for the different copies of a domain within the same gene; only data for high pLI EM genes are shown).

Finally, we observed that identical copies of a domain can be present in the same EM gene, and investigated whether each of those copies exerts equal contribution to gene function. We found substantial within-gene variability in CCR local constraint among identical domains (Figure 3c and Supplementary Figure S1a). For example, there are 7 copies of the PHD-finger domain in *KMT2D*, with a CCR local constraint ranging from 0 to 67. This variability is also present in non EM-specific domains (Supplementary Figure S1b), and exhibits interesting patterns. For instance, all 3 copies of the AT-hook DNA binding domain in *SRCAP* have a CCR local constraint of 0, whereas in *BAZ2A* the 4 copies have CCR local constraint of 0, 0, 20 and 100, showing that high heterogeneity in CCR local constraint can be gene specific. In contrast, the CCR local constraint of copies of the C2H2-like Zinc Finger domain are relatively uniformly distributed between 0 and 100 within each member of the PRDM family (Supplementary Table 4)

### A large subset of the epigenetic machinery is co-expressed

To identify functional properties specific to variation-intolerant EM genes, we systematically explored the expression patterns of the whole group across a spectrum of adult tissues, using publicly available RNA-seq data (GTEx Consortium, 2015). We selected 28 tissues on the basis of sample size and diversity in physiological function. First, we discovered that virtually all EM genes are expressed in a non-tissue specific manner, similarly to what is observed for known housekeeping genes (Methods, Supplementary Figure S2a,b), with the exception of a small number that showed testis-specific expression (Supplementary Figure S2c). Hence, tissue-specificity cannot account for the differences in variation tolerance within the EM group; we also found that it cannot explain the high mutational constraint of EM genes vs TF genes and other genes, after restricting the pLI comparison to very broadly expressed genes from both groups (Methods, Supplementary Figure S3).

Then, given that the transcriptional programs operating within each cell need to be precisely orchestrated, we hypothesized that at least some of the EM genes would show coordinated expression. To test this, we constructed tissue-specific co-expression networks and determined modules of co-expressed genes using WGCNA (Zhang and Horvath, 2005) (Methods). We noticed that for all tissues, EM genes were grouped in a few large modules (median 2 modules across tissues, range 0-4), with a substantial number of genes not belonging to any module (singletons; median 106 singletons across tissues, range 9-270). We asked if the division of EM genes into genes belonging to large modules and genes being singletons was stable across tissues. Because these modules were estimated separately for each tissue, it is not obvious how to compare them across tissues, and modules are affected by noise resulting from differences in sample size, and other sources. To perform the comparison, we defined two genes to be module partners if they belonged to the same module in at least 10 tissues, stable module partners if they belonged to the same module in at least 14 tissues, and not module partners if they belonged to the same module in less than 10 tissues (light blue, orange, and dark gray squares respectively in cartoon Figure 4a). For each gene we computed the number of module partners, and then ordered EM genes according to this score (Figure 4b). We next collectively visualized the pairwise partnership statuses among EM genes in a symmetric matrix, keeping this ordering for both rows and columns (Figure 4c, blue). We observed a distinct clustering, with a set of genes which are predominately stable module partners with each other (Figure 4c, orange), a large set of genes with no module partners (Figure 4c, dark gray), and a transition between these two groups (Figure 4c). We noted that the transition occurs as the number of module partners increases, meaning that EM genes not only have more partners, but they are also stable partners with the majority of them.

**Figure 4.**
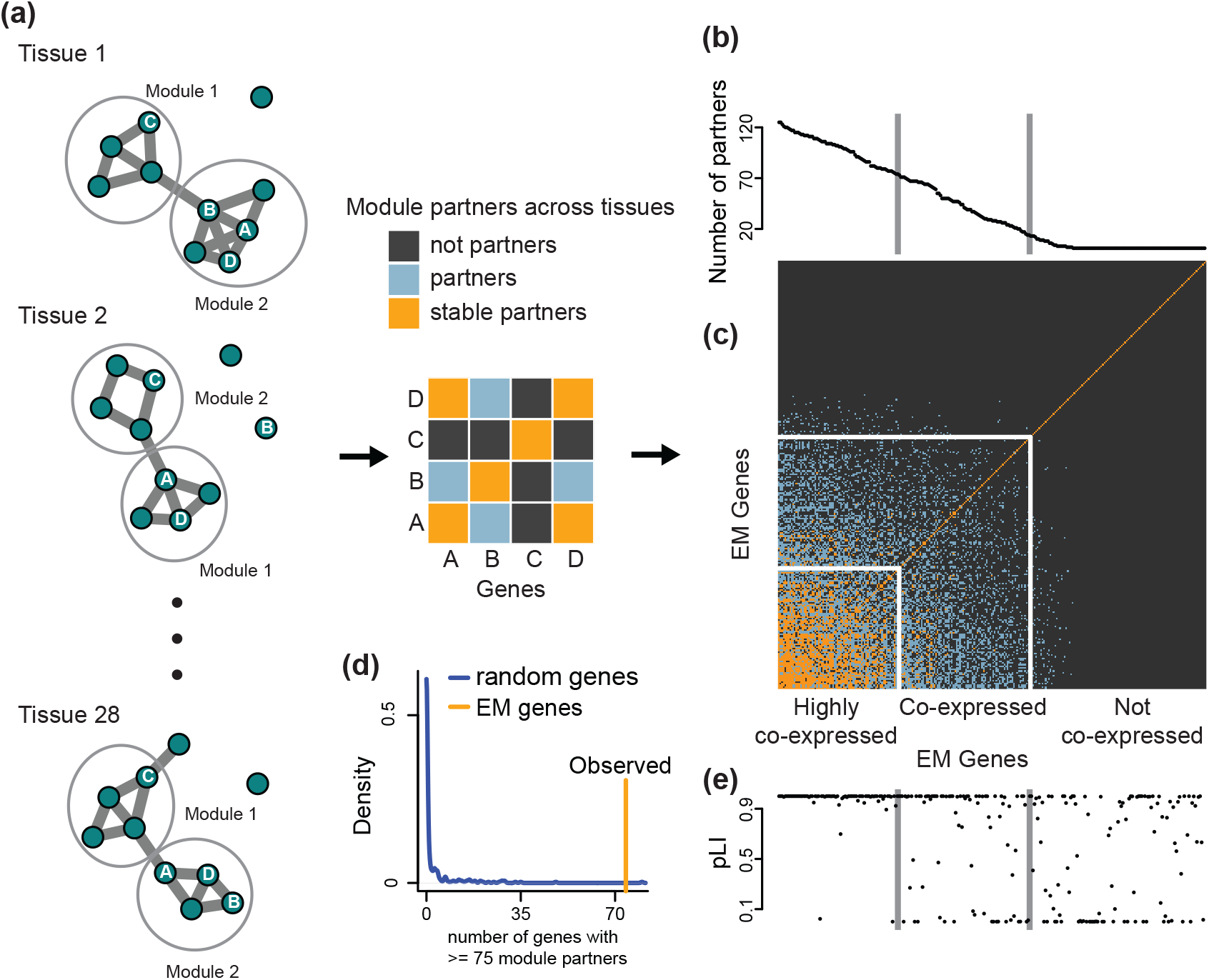
A large subset of the components of the epigenetic machinery exhibit unusually high levels of co-expression. **(a)** Schematic illustrating our definition and identification of module partners. WGCNA was used to construct tissue-specific co-expression networks and modules for 28 tissues profiled in GTEx. We determined if two EM genes were module partners (part of the same module in 10 – 14 tissues) or stable module partners (part of the same module in > 14 tissues). **(b, c)** The number of module partners for each EM gene and the module partner matrix, where rows and columns are ordered as in (b). We define 3 groups of EM genes, “highly co-expressed”, “co-expressed” and “not co-expressed” based on their number of module partners. **(d)** The pLI for each EM gene, ordered by the its number of module partners as in (b). **(e)** The size of the (highly) co-expressed group of EM genes compared to 300 draws of 270 random genes, where the random genes are selected to have a similar expression level across tissues compared to EM genes (Supplementary Figure S10).

We then divided EM genes into 3 groups: (1) a group of 74 genes with at least 75 module partners; we call this group of EM genes “highly co-expressed”, (2) a group of 83 genes with between 15 and 74 module partners; we call this group “co-expressed” and (3) a group of 113 genes with fewer than 15 module partners; we call this group “not coexpressed”. To assess the statistical significance of the size of these groups, we compared our results to those obtained with randomly chosen genes, and found that the groups of highly co-expressed as well as co-expressed EM genes are much larger than expected by chance (Figure 4e, Supplementary Figure S4a,b). We also established that our results are robust to the choice of cutoffs, the presence of sample outliers, and the exact network reconstruction method employed (Methods, Supplementary Figure S4c, Supplementary Figure S5). Finally, we note that our across-tissue co-expression analysis provides confidence that our findings are not driven by the cell-type heterogeneity present in these tissue samples.

### Highly co-expressed genes of the epigenetic machinery share common upstream regulators

To gain insights into the mechanistic basis of this co-expression, we first investigated the relative chromosomal positions of co-expressed EM genes. It is known that highly expressed genes tend to reside in chromosomal clusters in the human genome (Caron et al., 2001), and clustered genes are often co-expressed (Cohen et al., 2000). However, in this case we did not observe any notable clustering, despite the fact that EM genes have high levels of expression (Supplementary Figure S2d). The 74 highly co-expressed, and the 83 co-expressed EM genes were approximately uniformly distributed across chromosomes (chi-squared goodness of fit test, simulated *p* = 1 for both groups; Supplementary Figure S6; Supplementary Material).

We then asked if EM genes share upstream regulators, with the hypothesis that shared regulation induces co-expression. To answer this, we need robust data on the binding of many regulatory factors within a single cell type. For this reason, and since our previous results suggest that the co-expression is largely independent of cellular context in adults, we first focused on K562 cells, where ENCODE has produced good-quality data for 330 factors, more than any other cell line (Methods). We tested each of these factors for enriched binding at the promoters of the highly co-expressed EM genes relative to those of the non co-expressed EM genes, and found that 53 factors exhibit at least 2-fold enrichment with a p-value less than 0.05, in contrast to what is observed for randomly chosen genes, or after permuting the labels of EM genes (Supplementary Figure S7b and c, *p* = 0.02 and *p* = 5 · 10^−4^ respectively; Methods). Note that the direction of effect is consistent with our hypothesis: there is only one factor enriched in the non co-expressed group compared to the highly co-expressed group.

We next leveraged data from the Roadmap Epigenomics consortium (Roadmap Epigenomics Consortium et al., 2015) to obtain insight into fetal tissues, which are not represented in the GTEx dataset. Due to the lack of ChIP data on transcription factors in these samples, we used digital genomic footprinting data in fetal brain samples, and scanned the footprints that resided in EM promoters for known TF motifs (Methods) (Grant, Bailey, and Noble, 2011; Kulakovskiy et al., 2018); this approach has been proven useful in dissecting the architecture of gene regulatory networks (Neph et al., 2012). As for K562, we observed that the highly co-expressed EM genes are enriched for binding of common TFs compared to the non co-expressed group more than what would be expected by chance (Supplementary Figure S7d, *p* = 0.03; Methods).

Finally, we tested whether the co-expression is associated with protein-protein interactions (PPIs) between EM gene products, using recent data on such interactions (Huttlin et al., 2017). As our definition of EM genes only includes these with catalytic or reading activity, and not genes encoding for accessory subunits of chromatin modifying complexes, the extent to which such interactions will occur is not *a priori* known. We did not observe increased frequency of interactions between the highly co-expressed versus the non coexpressed group, with both groups having very few PPIs (probability of a pair interacting is 0.0007 and 0.003 in the two groups respectively).

### The co-expressed subset is extremely intolerant to variation and enriched for genes causing neurological dysfunction

We subsequently examined the relationship between co-expression and mutational constraint. Examination of the pLI scores of the three co-expression groups revealed a very clear association (Figure 4e, Figure 5a), with almost all highly co-expressed genes being extremely intolerant to variation (percentage of genes with pLI > 0.9 is > 90%), and co-expressed genes exhibiting intermediate intolerance.

**Figure 5.**
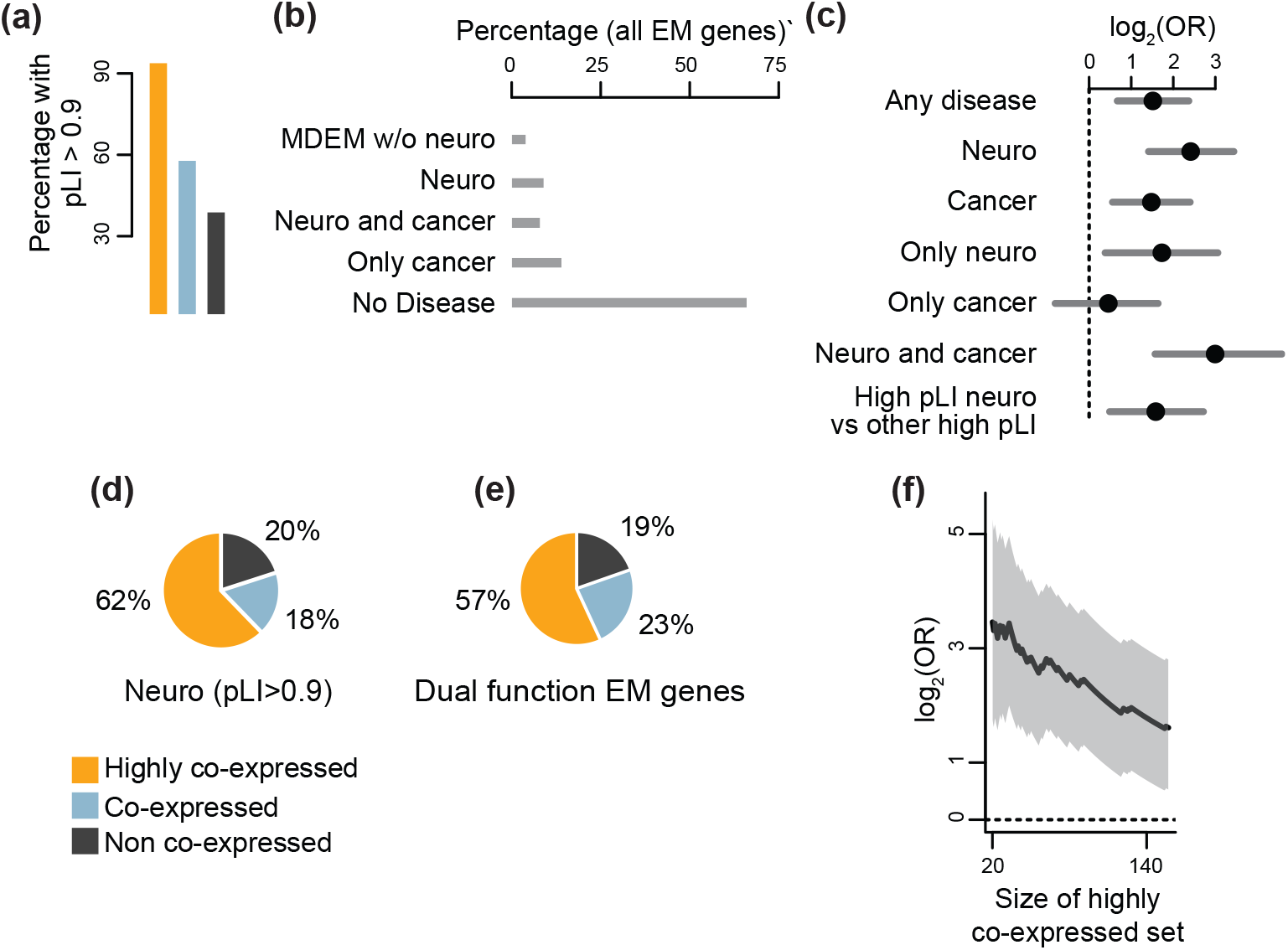
EM genes linked to disorders with neurological dysfunction demonstrate significant enrichment within the highly co-expressed category. **(a)** The percentage of EM genes with pLI > 0.9 in each of the co-expression categories. **(b)** The percentage of EM genes that are associated with different types of disease; individual disease categories are mutually exclusive. MDEM: Mendelian disorders of the epigenetic machinery, Neuro: includes autism, schizophrenia, developmental disorders, and MDEM whose phenotype includes dysfunction of the central nervous system (Methods). **(c)** Log odds ratios and 95% confidence intervals for enrichment of different subsets of EM genes in the highly co-expressed category. The dashed vertical line at 0 corresponds to statistical significance. **(d)** The percentage of EM genes that are associated with neurological dysfunction and have pLI > 0.9 in each of the co-expression categories. **(e)** The percentage of dual function EM genes in each of the co-expression categories. **(f)** Odds ratio (black line) and 95% confidence interval (shaded area) for enrichment of EM genes associated with neurological dysfunction in the highly co-expressed group, as a function of the size of the highly co-expressed group. For all sizes, the comparison was performed against the not co-expressed group.

We next asked whether, in addition to being very constrained, co-expressed EM genes are also preferentially associated with specific disease phenotypes. To perform this analysis, we first used our full list of EM genes to obtain a comprehensive picture of those links to disease. We examined associations with Mendelian disorders, cancer, and complex disorders (Methods). Consistent with previous observations (Bjornsson, 2015), we found that neurological dysfunction is a very prevalent phenotype within those diseases. Specifically, a total of 50 out of the 101 disease-associated EM genes genes have been previously describe to lead to neurological dysfunction (Figure 5b). Our analysis also yielded 64 EM genes associated with cancer (Figure 5b). We highlight a substantial overlap between those two groups: 24 of cancer associated EM genes are also associated with neurological dysfunction (Figure 5a), with dual function EM genes showing extremely high enrichment in this category (Fisher’s test, *p* = 1.84 · 10^—8^, odds ratio = 13.3). We did not find any EM gene both associated with cancer and also causing a Mendelian disease, without neurological dysfunction being part of the disease phenotype.

EM genes associated with any one of those disease phenotypes were enriched within the highly co-expressed group (Figure 5c; Fisher’s test, *p* = 2 · 10^—4^, odds ratio = 2.9). We thus sought to examine whether this enrichment was driven by associations with particular disease categories. We found a marked enrichment for genes causing neurological dysfunction (Figure 5c; Fisher’s test, *p* = 3.64 · 10^−7^, odds ratio = 5.3). Genes implicated in cancer were also enriched, although less (Figure 5c; Fisher’s test, *p* = 0.001, odds ratio = 2.8). We next tried to disentangle the contributions stemming from the associations with cancer versus neurological dysfunction. To accomplish this, we partitioned the EM genes into genes only associated with neurological dysfunction, genes only associated with cancer, and genes associated with both. Genes associated with neurological dysfunction were still enriched (Figure 5c; Fisher’s test, *p* = 0.006, odds ratio = 3.3), while we did not observe significant enrichment for genes only associated with cancer (Figure 5c; Fisher’s test, *p* = 0.41, odds ratio = 1.4). Subsequently, we asked if this result was a consequence of the association between co-expression and pLI. To answer this, we restricted the analysis to EM genes with high pLI (> 0.9). We found that neurological dysfunction were still significantly enriched in the highly co-expressed category (Figure 5c, d; Fisher’s test, *p* = 0.003, odds ratio = 3). Notably, genes associated with both neurological dysfunction and cancer were particularly highly enriched in the highly co-expressed group (Figure 5c; Fisher’s test, *p* = 4.04 · 10^−6^, odds ratio = 8); this was partially driven by the enrichment of dual function genes (Figure 5e; Fisher’s test, *p* = 9.16 · 10^−7^, odds ratio = 5.1). Finally, we examined the impact of the definition of the highly co-expressed group on the strength of the enrichment by varying the co-expression cutoff, while using the set of non-co-expressed EM genes as a control. As expected, this showed that the enrichment increased as the stringency of our cutoff increased (Figure 5c).

To further support our observed association between EM co-expression and neurological dysfunction, we asked whether the 16 TFs which show enriched binding at the promoters of the highly co-expressed group in fetal brain are associated with abnormal brain phenotypes. First, we found that 7 of the 16 have an associated syndrome in OMIM; for 4 out of 7, neurological dysfunction is part of the phenotype. Then, we used the Mouse Genome Database (Blake et al., 2017) to assess the phenotypes that result from genetic disruption of the mouse orthologues of these 16 factors. Phenotypes are available for 12 of the 16, and there is nervous system involvement in 8 out of 12. Collectively, there are 12 TFs for which there are phenotypic associations in mice or humans, and 9 of them are associated with neurological dysfunction in at least one of these organisms, independently supporting the association between EM co-expression and mammalian brain function (Supplementary Table 8).

## Discussion

We have performed a systematic investigation of all human genes encoding for epigenetic writers/erasers/remodelers/readers (EM genes). This enables us to make three basic contributions. First, we identify 103 novel disease candidates within this class of genes. Second, we provide strong evidence that chromatin state dysregulation is the most likely cause of the disease phenotypes. Third, we discover that co-expression distinguishes a large subset of EM genes that are both extremely variation-intolerant and, independently, enriched for genes causing neurological dysfunction.

We note however that our pLI-based approach for disease gene identification, while unbiased, cannot discriminate between genes that cause severe pediatric disease versus genes that lead to lethality at the embryonic stage. Additionally, while our local mutational constraint analysis argues that the epigenetic function itself is the primary driver of this intolerance to variation, some EM proteins may participate in biochemical events that affect other, non-chromatin related cellular functions (Spange et al., 2009; Biggar and Li, 2015). The importance of non-histone protein methylation and acetylation for signal transduction pathways and other molecular activities is not well understood, although there are examples of established functional relevance, such as cases of Cornelia de Lange syndrome caused by defective deacetylation of SMC3, a subunit of the cohesin complex (Deardorff et al., 2012). Further elucidation of the pathogenesis of cohesinopathies and related disorders may yield more insights into this issue.

Our most unexpected finding is that, among these 295 EM genes, we detected a subset of 74 that are highly co-expressed within tissues, as well as 83 others with an intermediate level of co-expression. The sizes of the two groups and the exact cutoff separating them might be refined with future interrogation of more tissues/cell types, and increases in sample size, but we anticipate the rank ordering of EM genes with respect to their partners to remain accurate. Surprisingly, this co-expression appears to unite three seemingly independent properties of the machinery: variation-intolerance, association with neurological dysfunction (even after conditioning on haploinsufficiency), and dual function (enzymatic activity combined with reading function). From a functional standpoint, the clear relationship between co-expression and mutational constraint indicates that the former potentially plays a role in homeostasis and disease predilection. It also suggests a basis for the observed dosage sensitivity, a counterintuitive result given that many EM genes are enzymes, and enzymes are usually haplosufficient (Jimenez-Sanchez, Childs, and Valle, 2001). For co-expressed enzymes however, a reduction of the normal amount of protein product present would not be tolerated, since it would compromise the coordinated expression of the module. Given the strong signal for enrichment of genes causing neurological dysfunction, it becomes tempting to speculate that the co-expression might be especially relevant to brain development and function; future examination of EM gene expression during fetal and early childhood development will likely yield profound insights into this. Finally, prioritization of highly co-expressed EM genes might not only aid in the discovery of new pathogenic variants disrupting the epigenetic machinery, but also provide a starting point for the interpretation of the functional consequences of those variants, particularly in the context of neurological dysfunction.

With respect to the underlying mechanistic basis of the co-expression, one way that such co-regulation could be achieved is with shared upstream regulators. Our data on regulatory factor binding at the promoters of EM genes support this possibility. However, a definitive answer to this will only be provided after further delineation of human regulatory circuits, with mapping of enhancer-promoter interactions in different cell-types. Our results also argue against the formation of multi-subunit complexes between the coexpressed EM gene products. Hence, it is possible that the need for co-expression arises not to regulate protein-protein interactions, but because imbalance of the epigenetic system could over time lead to major changes in open versus closed chromatin (Fahrner and Bjornsson, 2014).

In summary, our data provide the first evidence of widespread co-expression of epigenetic regulators, and link this novel phenomenon to both variation intolerance and neurological dysfunction, thus opening an avenue to better understand the role of the human epigenetic machinery in health and disease.

## Methods

### The creation of an epigenetic regulator list

We used InterPro domain annotations as provided by the UniProt database (Hunter et al., 2009; UniProt Consortium, 2015), accessed in June 2016, to generate a list of proteins with at least one domain that classifies them as writers or erasers of histone lysine methylation (Dillon et al., 2005; Shi, 2007), writers or erasers of histone lysine acetylation (Marmorstein and Zhou, 2014; Seto and Yoshida, 2014), readers of the two aforementioned histone modifications (Musselman et al., 2012), and readers of methylated and unmethylated CpG dinucleotides (Lee, Voo, and Skalnik, 2001; Jørgensen and Bird, 2002). A full list of all the domains used along with the corresponding InterPro IDs is provided in Supplementary Table 1. Additionally, we included the known human DNA methyltransferases and demethylases (Weissman, Naidu, and Bjornsson, 2014), as well as the catalytic subunits of the known human chromatin remodeling complexes (Clapier and Cairns, 2009). We also categorized all proteins belonging to the CHD family as chromatin remodelers (Marfella and Imbalzano, 2007), and we included the two histone lysine demethylases that do not harbor the JmjC domain, KDM1A and KDM1B (Shi, 2007). After manual curation, we excluded the proteins COIL, MSL3P1, ASH2L, PHF24, and VPRBP. We did not include the atypical histone lysine methyltransferase DOT1L, and we did not classify transcription factors whose recognition motifs include methylated CpG dinucleotides as DNA methylation readers. Proteins containing Ankyrin repeats were classified as histone methylation readers, provided they were first included as members of the epigenetic machinery based on the domains in Supplementary Table 1. We also note the case of the PHD finger domain: it was generally classified as a histone methylation reader domain, with the exception of 5 proteins (DPF1,2, and 3, and KAT6A,B) that have a double PHD finger which, based on experimental evidence (Zeng et al., 2010; Huber et al., 2017) acts as a histone acetylation recognition mode. We only included UniProt entries that have been manually annotated and reviewed by the database curators. The full list of all EM genes we used for our analyses along with several of their features, is given in Supplementary Table 2.

We note that while our categorization of EM-specific domains only includes domains which have some catalytic or reading function, there are domains which did not label as EM-specific, but are exclusively or almost exclusively found in EM genes. Two such examples are the pre-SET domain (present in 7 proteins, all of which are HMTs) and the post-SET domain (present in 16 proteins, 15 of those are HMTs and the remaining protein has 8 domains, all of which are post-SET domains).

### Epigenetic regulators with disease associations

With respect to Mendelian disease associations, we included disorders with a phenotype mapping key equal to 3 (indicating sufficient evidence to ascribe causality for a particular gene) in OMIM (https://omim.org/), as accessed in June 2016. We determined which of those syndromes involved dysfunction of the central nervous system based on the corresponding clinical synopses in OMIM. We also labeled the following genes as associated with neurological dysfunction: 1) genes that have been associated with Autism at a false discovery rate of 0.1 (De Rubeis et al., 2014) (those included the three genes later firmly associated with developmental disorders in Faundes et al. (2018)), 2) the top 15 % genes implicated in Schizophrenia (as ranked by their residual variation intolerance Score in McCarthy et al. (2014)), 3) *SETD1A*(Singh et al., 2016), 4) *KMT2B* (Zech et al., 2016), 5) genes that lacked previous associations with developmental disorders but achieved genome-wide significance in the DDD study (Deciphering Developmental Disorders Study, 2017), and examined the overlap with our set. Within our 103 novel disease candidates, we did not include candidate genes provided in Faundes et al. (2018).

Regarding associations with cancer, we first identified EM genes potentially functioning as cancer drivers using: 1) a list of 260 significantly mutated cancer genes, derived from data spanning 21 tumor types (Lawrence et al., 2014), and 2) genes that were predicted to be drivers by at least one of the top 3 performing methods in Tokheim et al. (2016). Both of the above studies evaluated genes based on point mutations and small insertions/deletions. We then also included other EM genes that have been reported to be involved in cancer, harboring either point mutations/small indels, or structural rearrangements (Shen and Laird, 2013; Feinberg, Koldobskiy, and Göndör, 2016). Collectively, we refer to all those EM genes as “cancer associated”.

All of the above disease associations are provided in Supplementary Table 2.

### Variation tolerance analysis

pLI scores for heterozygous loss of function constraint were downloaded from the ExAC database (Lek et al., 2016). When comparing the pLI distributions of different classes of genes, we excluded genes encoded on the X and Y chromosomes. PhastCons scores (Siepel et al., 2005) across 100 species were obtained for nucleotides using the GenomicScores and phastCons100way.UCSC.hg19 R packages. To calculate a gene level PhastCons score, we retrieved the hg19 genomic coordinates of all exons using the biomaRt (Durinck et al., 2005) R package (based on the same transcript ID as the one used in ExAC), and then used the function scores to get a PhastCons score for each exon (excluding the 5’ and 3’ UTRs); this score is the average PhastCons over the nucleotides in the exon. Subsequently, we calculated a weighted averaged over the exons (with the weights corresponding to length in bp) to derive a unique PhastCons score for each gene.

### CCR local constraint score

The CCR model (Havrilla et al., 2017) identifies regions of the genome without any missense or loss of function mutations in gnomAD (Lek et al., 2016). Each region devoid of mutations is assigned a CCR percentile score; the greater the difference between the observed and expected coverage-weighted length for regions with similar CpG density, the higher the constraint. As a result, the CCR model extends single gene-wide estimates of constraint to identify sub-regions within genes that exhibit “local constraint”. We mapped each protein domain to the genome, using the Pbase package. We defined the CCR local constraint for a domain to be the percentage of bases in the domain residing in a genomic region with a CCR percentile score above 90. The CCR local constraint score for protein domains in EM genes is included in Supplementary Table 4.

### GTEx data

RNA-seq data from 28 tissues (Supplementary Table 6), from 449 individuals were downloaded from the GTEx portal, release V6p. Those 28 tissues were selected based on differences in physiological function. Our goal was to obtain as representative a picture of human physiology as possible, but, since we ultimately performed across-tissue analyses, we sought to avoid the inclusion of tissues whose presence could introduce similarities between genes that would confound our tissue-specificity and co-expression analyses (see sections below). As an example, we only included samples from subcutaneous, and not from visceral adipose tissue.

For expression pattern analysis, we downloaded the raw RPKM data as provided in the GTEx portal. For co-expression analysis, we downloaded the gene-level count table and transformed to the log_2_(RPM + 1) scale (scaled to 10^7^ counts per sample instead of 10^6^). In this dataset, 5 EM genes were not available, leaving us with 290 for analysis.

### Tissue specificity and expression level analysis

Using the GTEx data described above, we calculated tissue-specificity scores for Figure S2 as previously described (Cabili et al., 2011). Computations were done using the functions `makeprobs()` and `JSdistFromP()` from the cummeRbund package (Trapnell et al., 2012). Figure S2b depicts the tissue-specificity scores of EM genes vs those of 30 genes encoding for TCA cycle related proteins (Supplementary Table 5). To confirm that our findings were not driven by unwanted variation, we repeated our analysis after correcting for RIN as well as surrogate variables (SVs) (Leek and Storey, 2007; Leek and Storey, 2008). In particular, using (log_2_ (RPKM + 1)) values, we estimated the SVs using the function `sva()`, from the SVA R package (Leek, Johnson, et al., 2012), while protecting for the tissue effect and including RIN as a known confounder. This resulted in 182 significant SVs, which we then used along with RIN to obtain the corrected expression values using the function `removeBatchEffect()` in the limma R package (Ritchie et al., 2015). Subsequently, negative values in the expression matrix were replaced by zeros, and genes with uniformly low values (< 0.01 in all samples) were removed. As depicted in Supplementary Figure S8), the results were essentially the same as those obtained with our original analysis. It should be mentioned however, that in situations where there is severe confounding of the factor of interest with some batch effect (for example, if samples from a particular tissue were all processed differently than others), correcting for surrogate variables cannot disentangle desired from undesired variation.

### Co-expression analysis

Using the GTEx data described above, we estimated tissue-specific networks and modules using the following approach. First, for each tissue, we only included genes where the corresponding tissue-specific median expression (median(log_2_(RPKM +1))) was greater than zero. Then, prior to network construction, we preprocessed the expression data to remove unwanted variation, since it is known that it can confound the estimation of pairwise correlation coefficients between genes (Freytag et al., 2015; Parsana et al., 2017). To achieve this, for each tissue, we standardized the expression matrix (containing (log2(RPM + 1)) values) to have mean 0 and variance 1 across every gene, and removed the 4 leading principal components from this matrix by regressing on the PCs and then reconstructing a new matrix with the regression residuals, using the function `removePrincipalComponents()` in the WGCNA package (Zhang and Horvath, 2005; Langfelder and Horvath, 2008). This has been shown to remove unwanted variation for co-expression analysis (Parsana et al., 2017). In Supplementary Figure S9 we depict the impact of doing this on the distribution of pairwise correlations across (1) 2000 randomly selected genes and (2) 80 genes encoding for the protein component of the ribosome (Supplementary Table 7), following ideas from Freytag et al. (2015). We expect random genes to be uncorrelated (negative controls) whereas we expect genes encoding for ribosomal proteins to be highly co-expressed (positive controls). To avoid over-fitting, we removed the same number of principal components from all tissues.

We then proceeded to perform tissue-specific network construction. For each tissue, we estimated the soft thresholding power using the entire expression matrix; we chose the first value for which the network was characterized by an approximately scale free topology, following standard WGCNA guidelines. To ensure that we can ultimately make comparisons across the 28 tissues, we next selected genes that have some minimal expression in all of them. Specifically, we required that the tissue-specific median expression (median(log_2_ (RPKM + 1))) was greater than zero in *all* of the 28 tissues. This gave us a set of 14872 genes. With respect to EM genes, this requirement selects 270 EM genes out of 295; in addition to the 5 of the 295 EM genes are not present in GTEx (see above), it excludes the 11 EM genes which are testis-specific (testis-specificity score > 0.5), 7 genes that are either not expressed or expressed at very low levels in those tissues, and 2 genes (*DPF1* and *PHF21B*) that are expressed at a considerable level in more than 1 tissue but are very lowly expressed in some other tissues (*DPF1* was especially expressed in cerebral cortex and cerebellum, but this was not as pronounced after correcting for RIN and surrogate variables). Subsequently, using only those 270 EM genes, we built unsigned tissue-specific networks and identified modules, by performing hierarchical clustering using the function `cutTreeDynamic()`, with the dissimilarity measure based on the topological overlap matrix. We set the parameters `minClusterSize` and `deepSplit` equal to 15 and 2, respectively. Modules were merged when the correlation between the corresponding module eigengenes was 0.8 or greater. Any parameters that are not mentioned were left at their default values.

To derive the reference distribution of the number of highly co-expressed and co-expressed genes (Figure 4d, Supplementary Figure S4), we performed all of the steps described above, but, instead of EM genes, with 270 randomly selected genes; this was repeated 300 times. Because we observed that EM genes which belong to either the highly co-expressed or co-expressed group have higher expression across tissues than EM genes which are not co-expressed (Supplementary Figure S10a), each time the random genes were sampled from the population of genes whose median expression level (median(log2 (RPKM + 1))) was: (1) at least 0.5 in more than half of the tissues (11963 genes total), or (2) at least 3 in more than half of the tissues (5095 genes total). These two populations of genes have a similar expression level to the EM genes (group 1) or a considerably higher expression level compared to the EM genes (group 2) (see Supplementary Figure S10b and c).

We examined the robustness of our results to the choice of arbitrary cutoffs, and determined that our choice of cutoffs do not impact the statistical significance of our finding (Supplementary Figure S4). To ensure that our findings are not driven by the presence of outlier samples, we compared our result to those obtained when we randomly excluded samples from the network construction. In particular: 1) when there were more than 100 samples in a tissue, we randomly dropped half of them, 2) when there were between 50 and 100 samples in a tissue we randomly dropped 20 of them, and when there were less than 50 samples we randomly dropped 5 of them. After this subsampling had taken place, the subsequent steps were performed as described above. This procedure was repeated 300 times. Across the random subsets, we found a median of 64.5 out of the 74 originally identified highly co-expressed genes being classified as such again.

For the analysis where we construct tissue-specific networks by thresholding the correlation matrix, we selected genes and removed principal components as described above. We then estimated a tissue-specific threshold as the 99.8% percentile of the correlation matrix of 2,000 randomly chosen genes. The topology of the resulting network is sensitive in both directions to the exact cutoff. We then thresholded the correlation matrix of the 270 EM genes and computed the maximally connected component of the resulting network. We then identified EM genes that shared membership in this component with more than 75 other EM genes in more than 10 tissues. We see 71 such genes, 60 of which are part of the originally detected highly co-expressed group, and 11 which are part of the co-expressed group, thus largely recapitulating the result obtained with WGCNA. We then compared the size and average node degree of the maximally connected component to those of the maximally connected components of networks constructed from groups of 270 randomly chosen genes; each of the reference distributions was derived by sampling 300 times from the population of genes with a similar expression level to EM genes (i.e. median expression greater than 0.5 in more than half the tissues, Supplementary Figure S10). We found that EM gene networks had a significantly larger maximally connected component, whose average node degree was generally also larger (Supplementary Figure S5).

### Regulatory factor binding at EM gene promoters

For this analysis, we defined promoters as 10 kb sequences centered around the transcriptional start site. We used the ENCODE portal (http://encodeproject.org), to download TF ChIP-Seq data for the K562 cell line (we note here that those data provide information for transcription factors, as well as other regulators not strictly belonging to the TF group; for simplicity, in this section we will refer to all those factors as TFs). Following ENCODE guidelines, we selected experiments that showed reproducibility across replicates and had narrow peak calls (that is, whose output type was labeled as “optimal IDR thresholded peaks”). Then, we only kept experiments performed in the absence of any treatment on the cells. We also randomly discarded any duplicate experiments (that is, experiments performed on the same TF target, regardless of the exact antibody used). Subsequently, we used the recount R package (Collado-Torres, Nellore, Kammers, et al., 2017; Collado-Torres, Nellore, and Jaffe, 2017) to select genes expressed in K562 cells (Slavoff et al. (2013); study identifier in the sequence read archive: SRP010061); as in our co-expression analysis, we required that a gene had median *RPKM* > 0 across the 8 total samples. This yielded a total of 242 EM genes (72 highly co-expressed and 94 non co-expressed), 14355 other genes (excluding ribosomal protein genes), and 330 regulatory factors.

To test for enrichment of TF binding in the highly co-expressed versus the non co-expressed EM gene group, we first discarded any TFs that were binding at only 10 promoters or less, as those were unlikely to be driving the observed co-expression. We then formed a 2*x*2 table for each of the remaining 295 TFs, and performed Fisher’s exact test. To derive a null distribution we used the following two approaches. First, we initially split the set of 14355 other genes to a set of 9495 genes with a median *RPKM* > 1.2 and a set of 10821 genes with a median *RPKM* > 0.4, to match the expression levels of the highly co-expressed and non co-expressed EM genes respectively (Supplementary Figure S7a). Then, we randomly sampled from these two sets to create two groups, consisting of 72 and 94 members respectively. We discarded any TFs binding at 50 promoters or less and then, as before, we tested each of the remaining 320 TFs for enrichment. Supplementary Figure S7b depicts the null distribution of the number of TFs with an Odds Ratio > 2 (indicating at least a 2-fold enrichment in the 72-member group) and a p-value < 0.05, versus the observed number of TFs showing this enrichment in the highly co-expressed EM group. For the second approach, we randomly sampled EM genes and after sampling we arbitrarily created two groups, one with 72 members and another with 94. We then repeated the same procedure as before, and tested each TF for enrichment in the promoters of one group versus the other. Supplementary Figure S7c shows the resulting null distribution, as well as the actual observed value.

For the analysis using digital genomic footprinting data, we used AnnotationHub to download footprint coordinates in the fetal brain. We then intersected the footprints with the promoters of EM genes, and obtained the sequences of the resulting set of promoter footprints. We subsequently scanned each of those sequences for TF motifs provided by HOCOMOCO (Kulakovskiy et al., 2018), using FIMO (Grant, Bailey, and Noble, 2011). We kept motif occurrences with a q-value less than 0.1. As we did not have expression data for those cells, we discarded any genes whose promoter footprints did not have any TF motifs after this analysis. We then aggregated the results at the promoter level, and we discarded TFs binding at 10 promoters or less. Those filters resulted in 259 EM genes (74 highly co-expressed and 107 non co-expressed), and 165 TFs. Similarly to the ChIP-seq analysis above, we then tested each TF for enriched binding at the promoters of the highly co-expressed EM group using Fisher’s exact test, and we derived a null distribution by randomly sampling EM genes and arbitrarily creating two groups with sizes equal to 74 and 107, respectively. The resulting null distribution is shown in Supplementary Figure S7d.

### Enrichment of disease categories in the highly co-expressed group

For Figure 5b we formed 2×2 tables of EM genes used in the co-expression analysis. For the categories “any disease”, “neuro” and “ca”, all 270 such genes were included. For the categories “neuro (no ca)”, and “ca (no neuro)” all EM genes associated with both neurological dysfunction and cancer were excluded. In all cases compared EM genes in the highly co-expressed group to EM genes in either the co-expressed or the not co-expressed group (combined). For the “high pLI neuro vs. other high pLI” category we only included genes with a pLI greater than 0.9 (without excluding those on the sex chromosomes). For Figure 5c we compared EM genes in the highly co-expressed group (defined using different cutoffs) to EM genes in the not co-expressed group, keeping the latter reference group constant in all comparisons.

## Disclaimer

The content is solely the responsibility of the authors and does not necessarily represent the official views of the National Institutes of Health.

## Supplementary Materials

### Variation intolerance of EM genes encoded on the sex chromosomes

In our main analysis of loss-of-function variation intolerance, we focused on genes encoded on the autosomes. When we exclusively considered the X chromosome, we observed a similar picture; 16 out of the 18 X-linked EM genes have a pLI greater than 0.9. Using data from a recent study on X inactivation (Tukiainen et al., 2017), we found that all 3 EM genes that consistently escape X inactivation in different tissues have a pLI of 1. In contrast, only 31% of other X-linked genes have a pLI greater than 0.9 (median pLI = 0.65, median pLI for other genes that escape X inactivation = 0.41). With respect to the 2 out of 4 EM genes on the Y chromosome that are included in ExAC, *UTY* has an intermediate pLI of 0.63, while *KDM5D* is haplosufficient (pLI = 0.02).

### Across-species conservation of EM genes

We examined the across-species conservation of EM genes, and compared them to both TF genes and all other genes. We used PhastCons scores calculated across 100 species (Siepel et al., 2005), and aggregated these into a gene-level score (Methods). In this analysis, TF genes are less conserved than other genes (Wilcoxon rank sum test, *p* = 0.002, Figure 2c), however, EM PhastCons scores are significantly more skewed towards 1 (Wilcoxon rank sum test, *p* = 2.21 · 10^−12^ and *p* = 4.71 · 10^−10^, Figure 2c). In general, EM genes with high pLI are characterized by a higher PhastCons score (Figure 2d). We did however, identify four genes with high pLI, but PhastCons score less than 0.5 (gray shaded area in Figure 2d): the eraser-reader *KDM4E* (pLI = 0.76 and PhastCons score = 0.44), the erasers *KDM7A* and *HDAC5* (pLI = 1 for both, and PhastCons score = 0.50 and 0.49, respectively), and the reader *BAZ2B* (pLI = 1 and PhastCons score = 0.39). Finally, we note the existence of a substantial number of EM genes with high PhastCons score and low pLI (Figure 2d).

### Tissue specificity and expression levels of EM genes

The intolerance to variation suggests that for most of the EM genes, the loss of even a single copy is incompatible with a healthy organismal state. An important question then, is the identification of the tissues and cell types through which this detrimental effect is mediated. There are two primary reasons why one might speculate that some EM genes have tissue-specific expression. First, the composition of the machinery suggests the existence of functionally redundant components (for instance, there are 116 histone methylation readers with no enzymatic or other reading activity). This redundancy could be explained if different components with the same role were specific to different tissues. Second, there is some evidence suggesting that TF binding to their target sites requires, at least in some cases, an already permissive chromatin state Guertin and Lis, 2013. This could imply that there are certain EM components expressed in a cell-type specific fashion, that thereby help generate the epigenomic landscapes that facilitate TF binding (Guertin and Lis, 2013; Quante and Bird, 2016). Given that specific genomic locations need to be marked, it has been postulated that this is achieved by EM genes with DNA-binding domains that differ from those encountered in classical TFs, yet confer some degree of sequence specificity (Guertin and Lis, 2013; Quante and Bird, 2016). Four domains described as putative candidates are the ARID DNA-binding domain, the AT-hook DNA-binding motif, the CxxC domain, and the C2H2-like Zinc finger, which we found to be present in 21 EM genes.

To gain insight into the question of tissue-specificity, we examined the expression patterns of EM genes across a spectrum of adult tissues, using RNA seq data generated by the GTEx consortium (GTEx Consortium, 2015). We selected 28 tissues on the basis of sample size and differences in physiological function, as this enabled us to obtain a comprehensive picture under diverse cellular conditions, and allowed us to avoid spurious specificity estimates arising from the high similarity between some tissues (e.g. subcutaneous and visceral adipose tissue). For each EM gene, we calculated an entropy based tissue-specificity score, as previously described (Cabili et al., 2011). This score reflects the degree to which a gene is highly specific for some tissue (score close to 1) or is expressed broadly across tissues (score close to 0). We discovered that the vast majority of EM genes are characterized by very low specificity, after comparing their scores to those of TF genes as well as other genes (Figure S2a). In fact, when we compared the specificity of EM genes to that of genes encoding for proteins involved in the tricarboxylic acid (TCA) cycle, a well known category of housekeeping genes, we found a very similar distribution (Figure S2b). The lack of tissue specificity characterizes EM genes with a DNA-binding domain that recognizes short motifs as well (median specificity score = 0.1), although we note that there also exist other genes that harbor those domains but do not fulfill the criteria for inclusion in our list of EM genes. This result however, raises the possibility that the increased dosage sensitivity of EM genes is due to their non-specific expression pattern. However, even after considering only highly non-specific genes (score less than 0.1), the enrichment of the 160 EM genes satisfying this criterion in the highly constrained category remains extremely pronounced (Supplementary Figure S3). This is true both when comparing to all of the 5249 other non-specific genes (Fisher’s test, *p* < 2.2 · 10^−16^, odds ratio = 5.6), as well as to the 232 non-specific TFs (Fisher’s test, *p* = 6.73 · 10^−9^, odds ratio = 3.5), showing that this constraint is not merely a consequence of the presence of EM genes in a greater number of cell types, but rather reflects their function.

We subsequently reasoned that our analysis might be masking the presence of genes specific for only a small subset of tissues, and we performed a detailed analysis of the specificity of EM genes separately for each tissue (Figure S2c). We observed that testis stands out as the only tissue for which a small number of EM genes show specific expression (Figure S2c), indicating its dependence on not only the general machinery that operates in all other tissues, but also on a distinct subset of components. This is in agreement with the existing view that testis is an outlier tissue with respect to its transcriptomic state (GTEx Consortium, 2015). The testis-specific EM genes include *PRDM9*, in accordance with its reported role in meiotic recombination (Hochwagen and Marais, 2010), as well as 10 other genes (Figure S2c), some of which (*TDRD1, RNF17, BRDT, PRDM14, MORC1*) possibly play roles in male germ cell differentiation and the repression of transposable elements in the germline (Chuma et al., 2006; Pan et al., 2005; Shang et al., 2007; Yamaji et al., 2008; Pastor et al., 2014), while the role of the others (*CDY2A, HDGFL1, PRDM13, PRDM7, TDRD15*) remains mostly unspecified. Three of those genes are also members of the PRDM family of histone methyltransferases, while the rest are all readers of histone methylation, with the exception of *BRDT*, a histone acetylation reader.

Finally, in all of the tissues analyzed, we observe that EM genes are highly expressed compared to TF genes, and other genes (Figure S2d). We also confirm that TF genes are expressed at low levels (Figure S2d), as was previously observed using microarray data (Vaquerizas et al., 2009). Furthermore, we find an association between pLI and expression level (Supplementary Figure S11), which could be attributed to the fact that a reduction of this high expression into half the normal amount is not tolerated. As expected, there are exceptions, namely TF genes or other genes that are expressed at equally high or higher levels than EM genes, as well as a median of 44.5 EM genes across tissues expressed at low levels, with a median RPKM always less than 1. Within the latter, 9 components show consistently low expression across all tissues, with 6 being testis-specific. Collectively, the above results indicate that the majority of human EM genes are active across a heterogeneous set of adult tissues. This suggests that other factors primarily maintain cell identity in those tissues, and it can help explain the observation that in most cases of Mendelian disorders of the epigenetic machinery, more organ systems are affected compared to other genetic disorders (Bjornsson, 2015).

### Spatial clustering of co-expressed EM genes

Given that highly expressed genes in the human genome tend to reside in chromosomal clusters (Caron et al., 2001), and taking into account that clustered genes are often co-expressed (Cohen et al., 2000), we investigated the relative chromosomal positions of co-expressed EM genes. We did not observe any notable clustering, with our 74 highly co-expressed, and 82 co-expressed EM genes being approximately uniformly distributed across chromosomes (chi-squared goodness of fit test, *p* = 1 for both groups), and only 9 and 6 pairs, respectively, having a within-pair chromosomal distance less than 1 megabase (Supplementary Figure S6). The single exception to this is *SETD1A* and *FBXL19*, which are separated by only 8 kb. To see if the co-expression of those two genes is driven by a bidirectional promoter we looked at whether they are encoded on opposite strands, and found this not to be the case.

## Supplementary Tables

We provide a number of Supplementary Data Files.

1. **Supplementary Table 1:** The EM-specific protein domains used to define the components of the epigenetic machinery.
2. **Supplementary Table 2:** The components of the epigenetic machinery. Each row corresponds to a gene, and has the following columns:

a. `Entry:` Entry ID in UniProt.
b. `Status:` Whether the entry has been reviewed by the database curators.
c. `Gene _name:` Gene name.
d. `Alt_Gene_names:` Alternative gene names.
e. `Chr:` Chromosome.
f. `Strand:` Strand.
g. `Start_Pos_hg38:` Start Position (in hg38 coordinates).
h. `End_Pos_hg3 8:` End Position (in hg38 coordinates).
i. `Protein.names:` Protein name (with alternative protein names in parentheses)
j. `Interpro.Domains:` InterPro domain IDs
k. `Epi_function:` Epigenetic function (e.g. “Writer”)
l. `Enzymatic.activity:` Enzymatic activity (HMT: histone methyltransferase, HAT: histone acetyltransferase, DNMT: DNA methyltransferase, HDM: histone demethylase, HDAC: histone deacetylase, DNM_ERASE: DNA demethylase, REMODEL: chromatin remodeler).
m. `Reading.activity:` Reading activity (HMR: histone methylation reader, HAR: histone acetylation reader, DNMR: DNA methylation reader, DNUMR: reader of unmethylated CpGs).
n. `TF_activity:` Whether the gene is also classified as a Transcription Factor.
o. `OMIM_Mendelian_phenotype_ (number of phenotype):` OMIMMendelian phenotypes with a phenotype mapping key equal to 3.
p. `Inheritance_of_mend_ph_ (number of phenotype):` the corresponding modes of inheritance for each of the Mendelian phenotypes.
q. `is.neuro.associated:` Whether the gene causes a Mendelian disorder whose phenotype includes neurological dysfunction, or has been implicated in Autism, Developmental Disorders, or Schizophrenia.
r. `is.ca.associated:` Whether the gene has been implicated in cancer.
s. `Gene_name_exac:` Gene name in the ExAC database,
t. `exac.pLI:` pLI score as provided by ExAC.
u. `exac.pRec:` pRec score as provided by ExAC.
v. `exac.pNull:` pNull score as provided by ExAC.
w. `exac.mis_z:` Missense Z score as provided by ExAC.
x. `Gene_name_gtex:` Gene name in the GTEx dataset.
y. `coexpression_status:` Whether the gene is highly co-expressed, co-expressed, or not co-expressed according to our analysis. “NA” indicates that it was not included in the co-expression analysis.
3. **Supplementary Table 3:** Novel EM disease candidate genes, along with their coexpression status.
4. **Supplementary Table 4:** Local CCR constraint for all domains (rows), for all genes included in the local constraint analysis.
5. **Supplementary Table 5:** The 30 genes encoding for TCA cycle related proteins.
6. **Supplementary Table 6:** GTEx tissues used for the analyses.
7. **Supplementary Table 7:** 80 genes encoding for protein components of the ribosome.
8. **Supplementary Table 8:** The Transcription Factors enriched for binding at the promoters of highly co-expressed EM genes, along with the enrichment statistics and their existing phenotype associations in human and mouse.

## Supplementary Figures

**Supplementary Figure S1.**
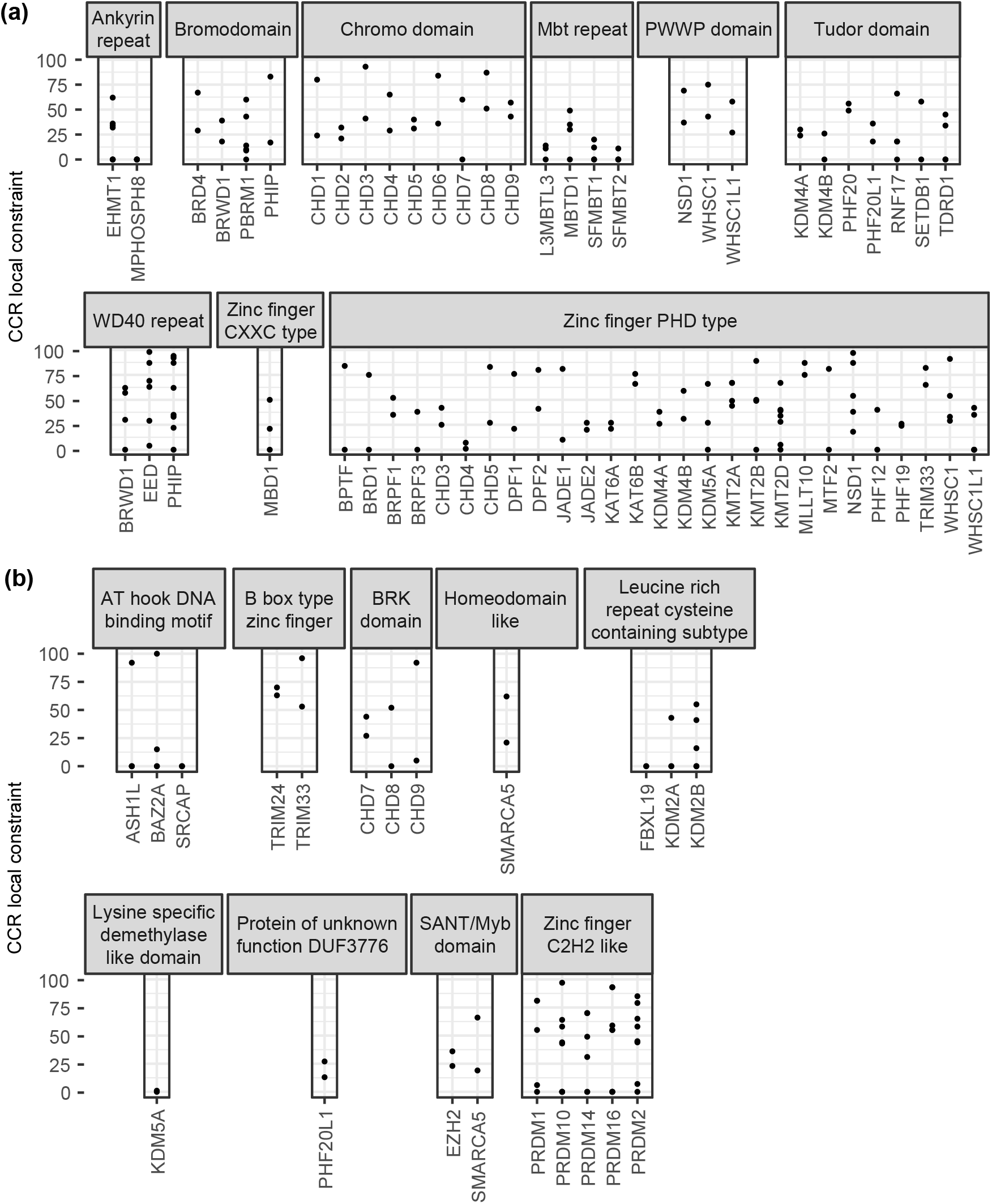
Identical copies of protein domains show within-gene variability in constraint. Each plot corresponds to a domain, and the points therein are the CCR local constraint scores for the different copies of the same domain within each gene. Only genes with more than one copy of the particular domain are shown. **(a)** EM-specific domains. **(b)** non EM-specific domains in EM genes.

**Supplementary Figure S2.**
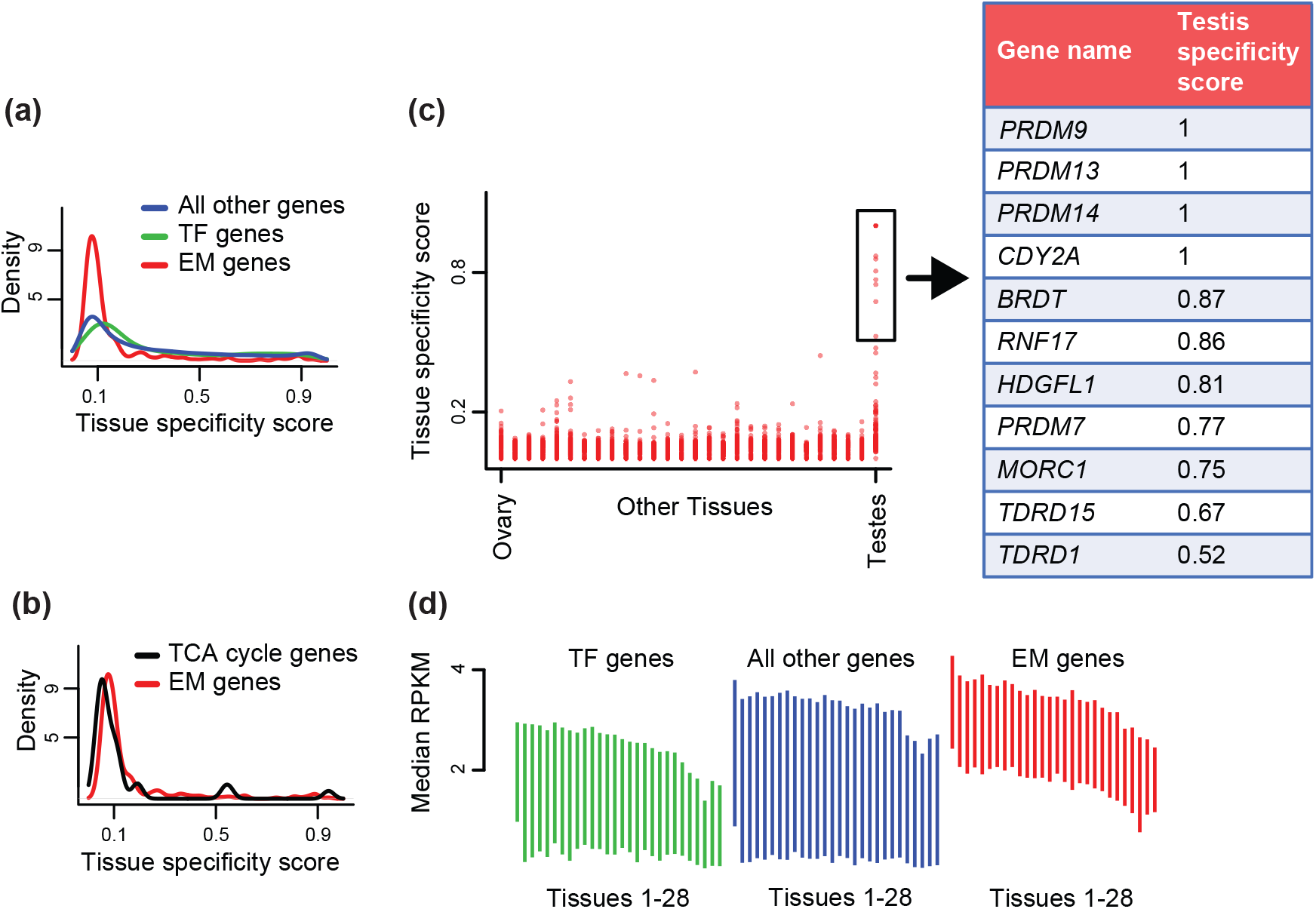
The components of the epigenetic machinery are expressed in a highly non tissue-specific manner and at high levels across tissues. **(a)** The distribution of the tissue-specificity score of EM genes (red curve) reveals their lack of tissue-specific expression, compared to TF genes (green curve), and all other genes (blue curve). **(b)** Comparison of the tissue-specificity of EM genes (red curve) with that of genes encoding for tricarboxylic acid (TCA) cycle related proteins (black curve) shows that EM genes exhibit comparable tissue-specificity to this class of well known housekeeping genes. **(c)** Testis is the sole tissue for which some EM genes have high specificity. **(d)** A comparison of expression levels of EM genes (red boxes) to those of TF genes (green boxes) and all other genes (blue boxes) shows their high relative expression. Each box shows the inter-quartile range of expression values, and tissues are ordered according to median expression for EM genes.

**Supplementary Figure S3.**
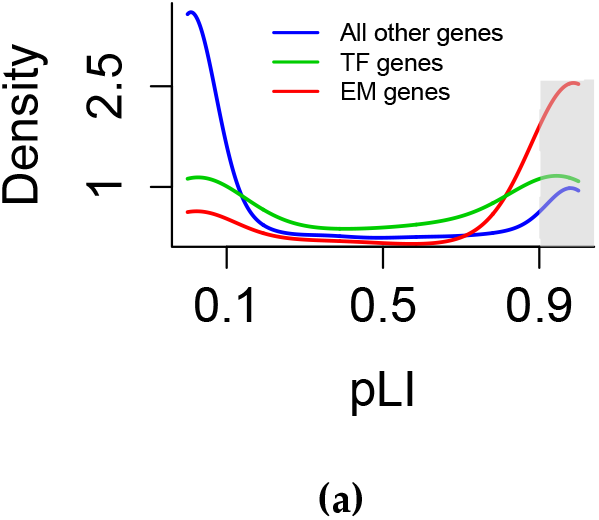
The pLI distributions of genes with low tissue-specificity. Density plots of pLI scores for genes with low tissue-specificity score (< 0.1) highlight that the enrichment of EM genes (red curve) in the highly intolerant category (pLI > 0.9, gray shaded area) compared to TF genes (green) and other genes (blue) remains very pronounced even after considering only broadly expressed genes.

**Supplementary Figure S4.**
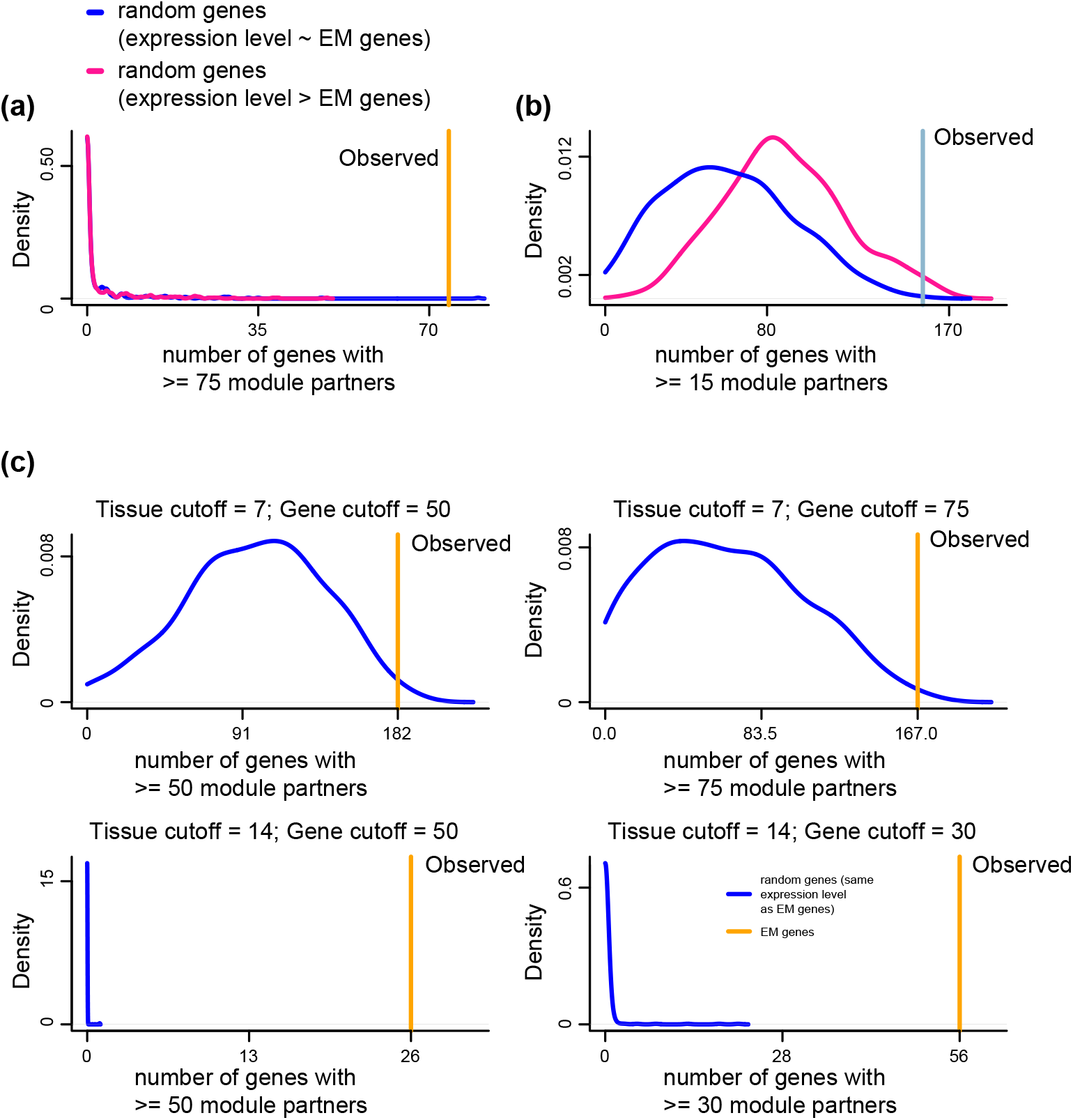
EM genes are highly co-expressed irrespective of arbitrary choices. We examine the sensitivity wrt. various cutoffs of the result that EM genes are highly co-expressed. **(a)** Like Figure 4d, but with an additional reference distribution where random genes are selected to have higher expression level than EM genes (Supplementary Figure S10). **(b)** Like Figure 4d but where we consider our observation that we have 157 EM genes which are either highly co-expressed or co-expressed. **(c)** Like Figure 4d, but for various choices of arbitrary cutoffs. Specifically we vary (1) in how many tissues two genes need to belong to the same module, to be considered partners (“tissue cutoff”, in the main text we use 10) and (2) how many module partners a gene needs to have, to be considered part of the highly co-expressed group (“gene cutoff”, in the main text we use 75).

**Supplementary Figure S5.**
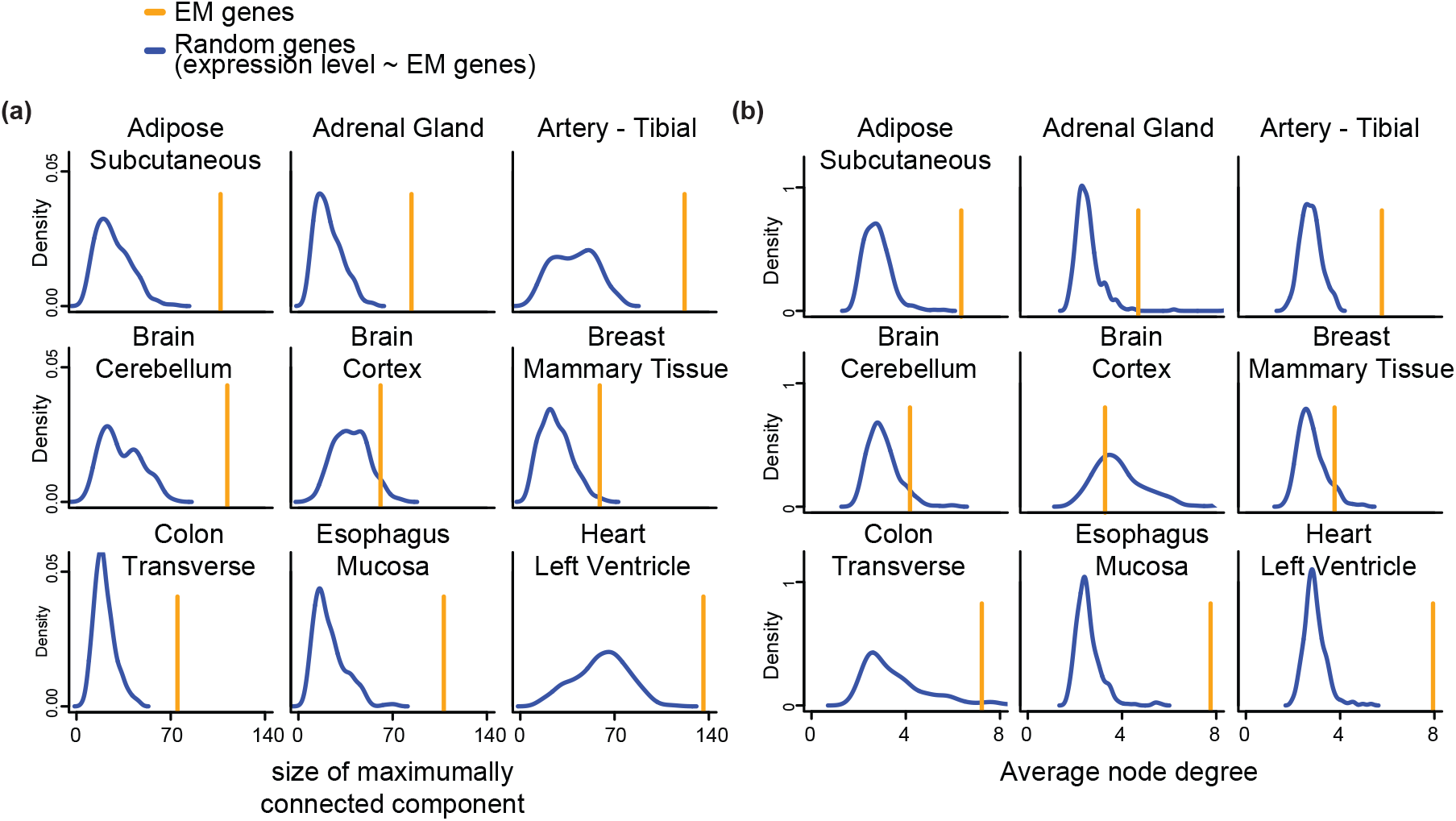
Size and average degree of the maximally connected component in different tissues. We estimated tissue-specific networks by thresholding the correlation matrix. For each tissue we computed the maximally connected component of the EM genes. As comparison we did the same for 300 random samples of 270 genes with a similar expression level to the EM genes. Depicted are results from 9/28 tissues (same tissues as in Supplementary Figure S9). **(a)** The size of the maximally connected component of the EM genes compared to the random samples. The component of the EM genes is significantly larger in 8 of the 9 tissues. **(b)** The average node degree within the maximally connected components. The EM genes are significantly more connected in 6 of the 9 tissues.

**Supplementary Figure S6.**
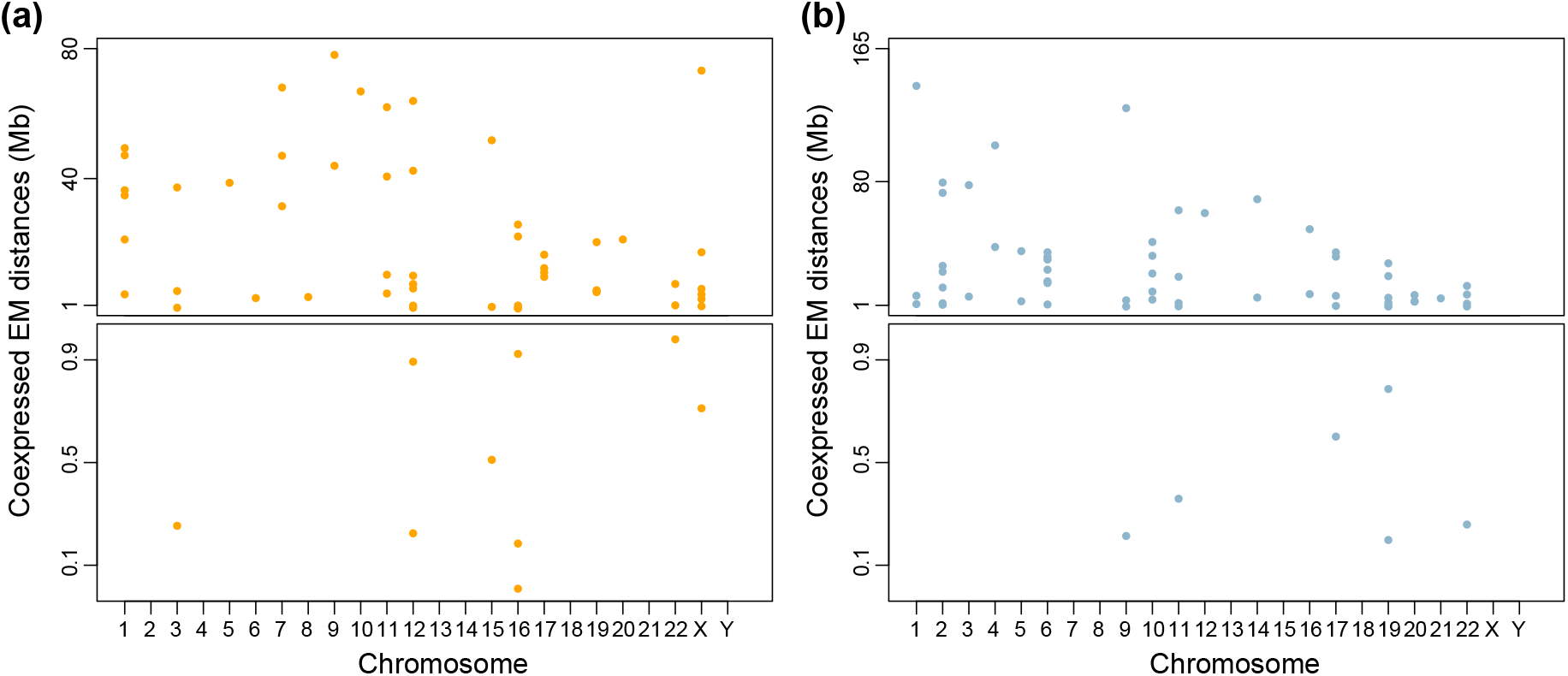
Co-expressed EM genes are not spatially clustered. The pairwise chromosomal distances (number of bp separating the transcription end site of a gene with the transcription start site of the most proximal downstream gene) between EM genes. Top panel are distances greater than 1 Mb and bottom panel less than 1 Mb. **(a)** Highly co-expressed EM genes. **(b)** Co-expressed EM genes.

**Supplementary Figure S7.**
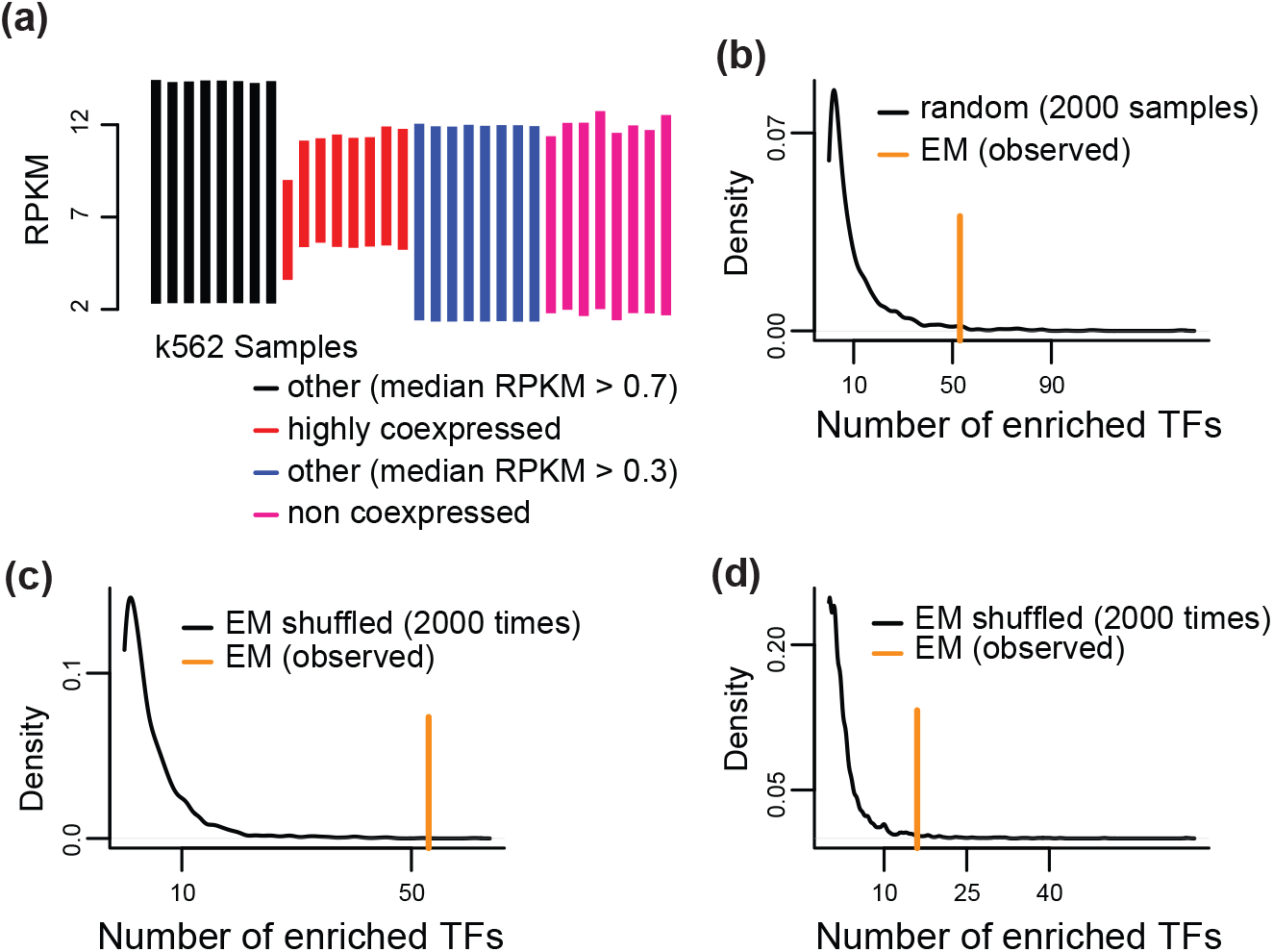
The promoters of the highly co-expressed EM genes are bound by common regulatory factors. **(a)** The distributions of expression levels in K562 cells for genes that were used to derive the reference distribution in **(b)**. **(b)** The distribution of the number of regulatory factors that show enriched binding (log_2_ OR > 1 and *p* < 0.05) at the promoters of the first group created after randomly sampling genes in K562 cells (black curve), versus the observed number of regulatory factors showing enriched binding at the promoters of highly co-expressed EM genes (orange vertical line). **(c)** The distribution of the number of regulatory factors that show enriched binding (log_2_ OR > 1 and *p* < 0.05) at the promoters of the first group created after shuffling the labels of EM genes in k562 cells (black curve), versus the observed number of regulatory factors showing enriched binding at the promoters of highly co-expressed EM genes (orange vertical line). **(d)** The distribution of the number of transcription factors that show enriched binding (log_2_ OR > 1 and *p* < 0.05) at the promoters of the first group created after shuffling the labels of EM genes in fetal brain (black curve), versus the observed number of regulatory factors showing enriched binding at the promoters of highly co-expressed EM genes (orange vertical line). For **(b)** and **(c)**, binding is measured by ChIP-Seq, whereas for **(d)** binding is measured by digital genomic footprinting (Methods).

**Supplementary Figure S8.**
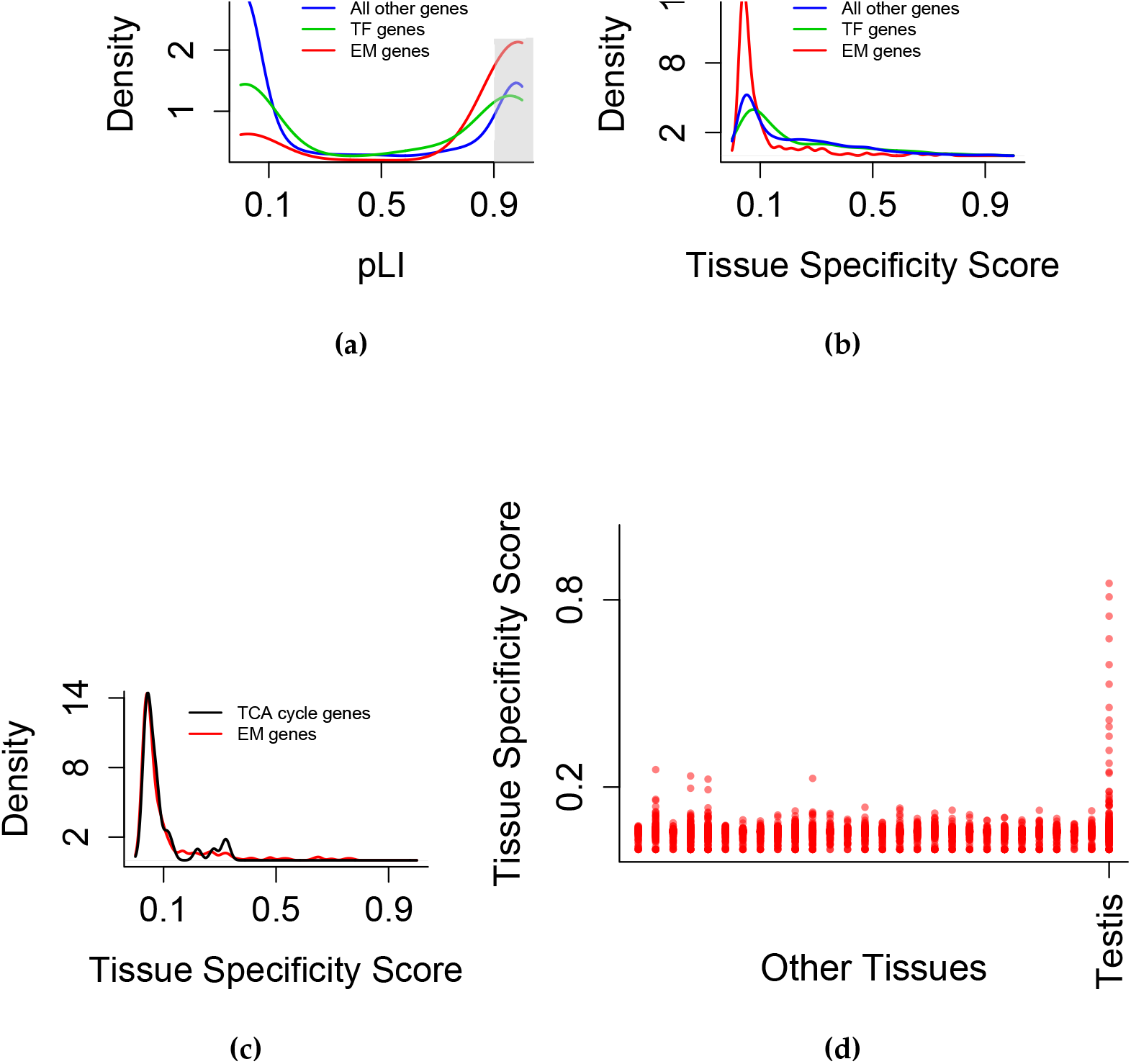
The lack of tissue-specificity of EM genes is not driven by unwanted variation. Results for the tissue-specificity analyses after correcting for RIN and surrogate variables (Methods). **(a)** Like Supplementary Figure S3. **(b)** Like Figure S2a. **(c)** Like Figure S2b. **(d)** Like Figure S2c.

**Supplementary Figure S9.**
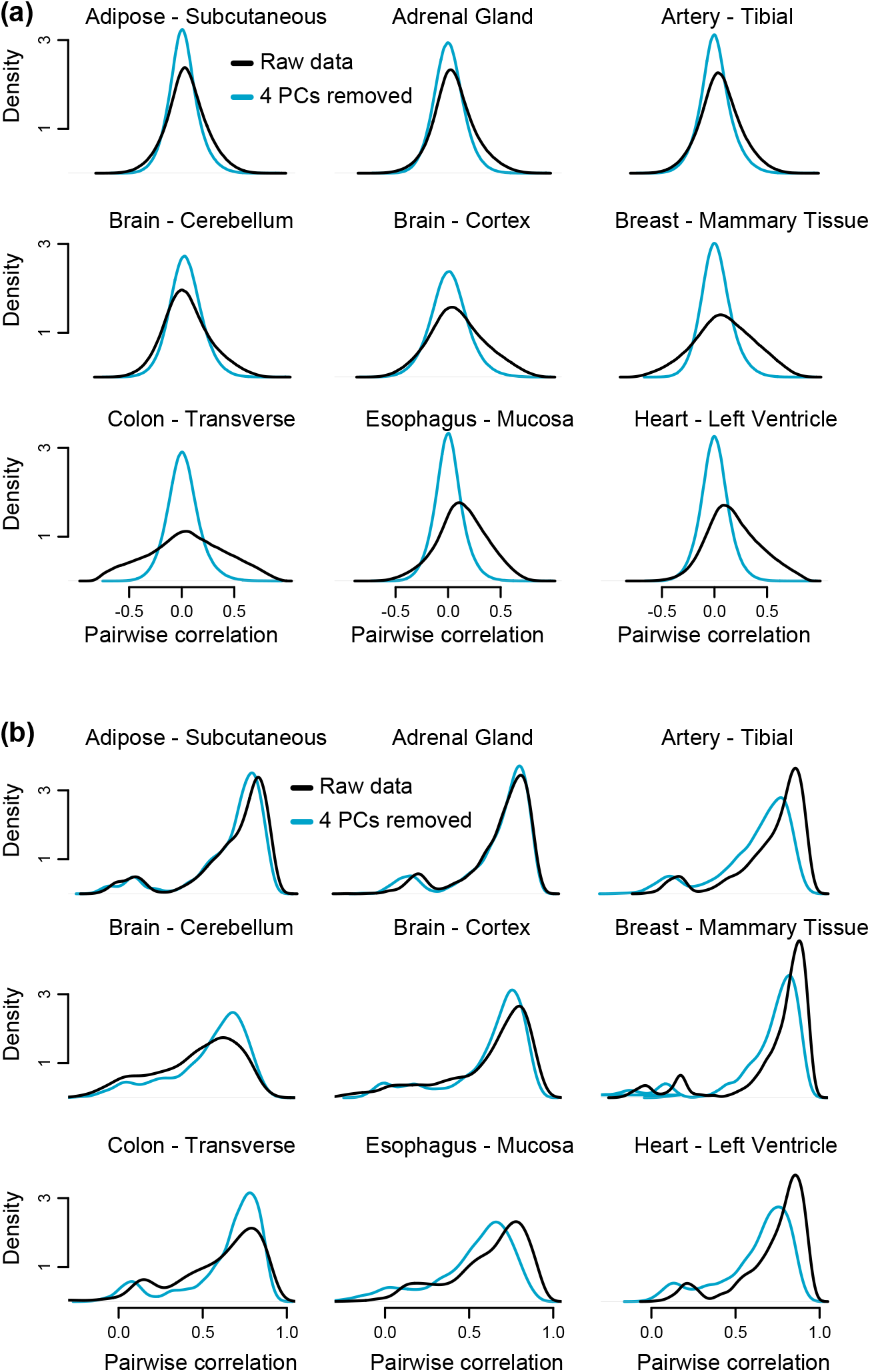
Removing noise in co-expression analysis by removing principal components. We remove unwanted variation in our co-expression analysis by removing 4 principal components from the expression matrix in all tissues. **(a)** The distribution of pairwise correlations between randomly sampled genes, serving as a negative control, for 9 out of the 28 tissues. **(b)** The distribution of pairwise correlations between 80 genes coding for ribosomal proteins, which serve as a positive control, for the same tissues as in (a).

**Supplementary Figure S10.**
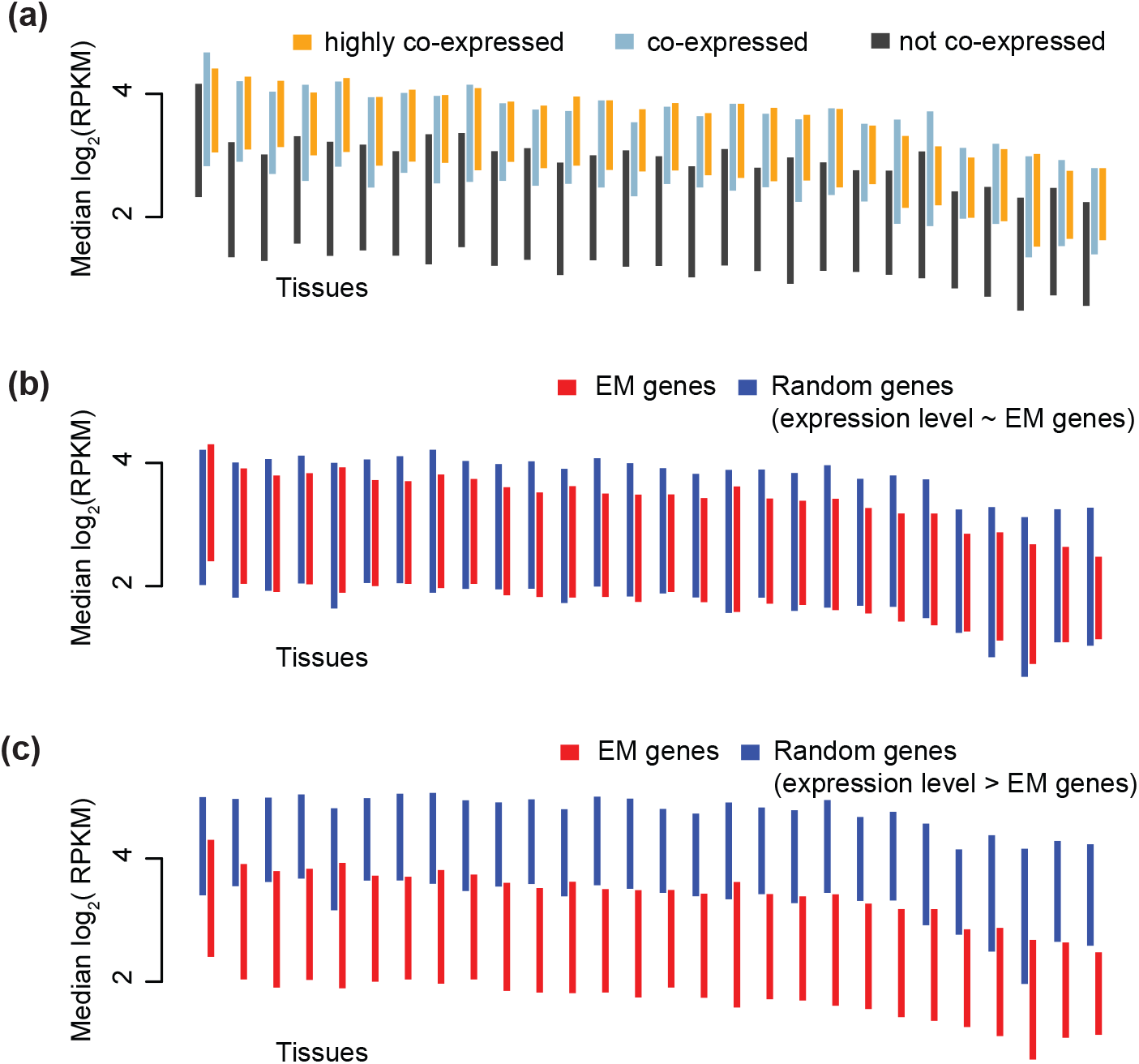
Expression levels of EM genes. Expression (log(RPKM + 1))) for various groups of genes. **(a)** We categorize EM genes into 3 groups based on co-expressed module patterns across tissues (Figure 4). EM genes which are highly co-expressed or co-expressed have a higher expression level than EM genes which are not co-expressed. The former two categories show similar expression levels. **(b)** The expression level of EM genes compared to 11963 genes where the median expression in each tissue is greater than 0.5 in more than half the tissues. The two groups of genes have similar expression level. We say these genes are similarly expressed to the EM genes. **(c)** The expression level of EM genes compared to 5095 genes where the median expression in each tissue is greater than 3 in more than half the tissues. The latter group of genes are expressed at higher levels than the EM genes.

**Supplementary Figure S11.**
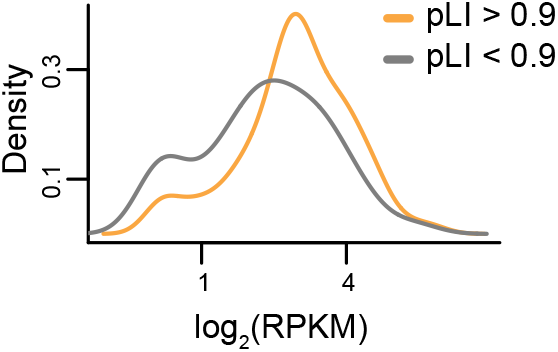
Highly constrained EM genes are also more highly expressed. Density plots of log 2 (RPKM) values for EM genes with pLI > 0.9 vs. those of EM genes with pLI < 0.9 show that the former exhibit higher expression levels.

